# Structures reveal how the Cas1-2/3 integrase captures, delivers, and integrates foreign DNA into CRISPR loci

**DOI:** 10.1101/2025.06.10.658980

**Authors:** William S. Henriques, Jarrett Bowman, Laina N. Hall, Colin C. Gauvin, Hui Wei, Huihui Kuang, Christina M. Zimanyi, Edward T. Eng, Andrew Santiago-Frangos, Blake Wiedenheft

## Abstract

Cas1 and Cas2 are the hallmark proteins of prokaryotic adaptive immunity. However, these two proteins are often fused to other proteins and the functional association of these fusions often remain poorly understood. Here we purify Cas1 and the Cas2/3 fusion protein from *Pseudomonas aeruginosa*. We determine multiple structures of the Cas1-2/3 complex at distinct stages of CRISPR adaptation. Collectively, these structures reveal a prominent, positively charged channel on one face of the integration complex that captures short fragments of foreign DNA. Foreign DNA binding triggers conformational changes in Cas2/3 that expose new DNA binding surfaces necessary for homing the DNA-bound integrase to specific CRISPR loci. The length of the foreign DNA substrate determines if Cas1-2/3 docks completely onto the CRISPR repeat to successfully catalyze two sequential transesterification reactions required for integration. Taken together, these structures clarify how the Cas1-2/3 proteins orchestrate foreign DNA capture, site-specific delivery, and integration of new DNA into the bacterial genome.

**GRAPHICAL ABSTRACT:** 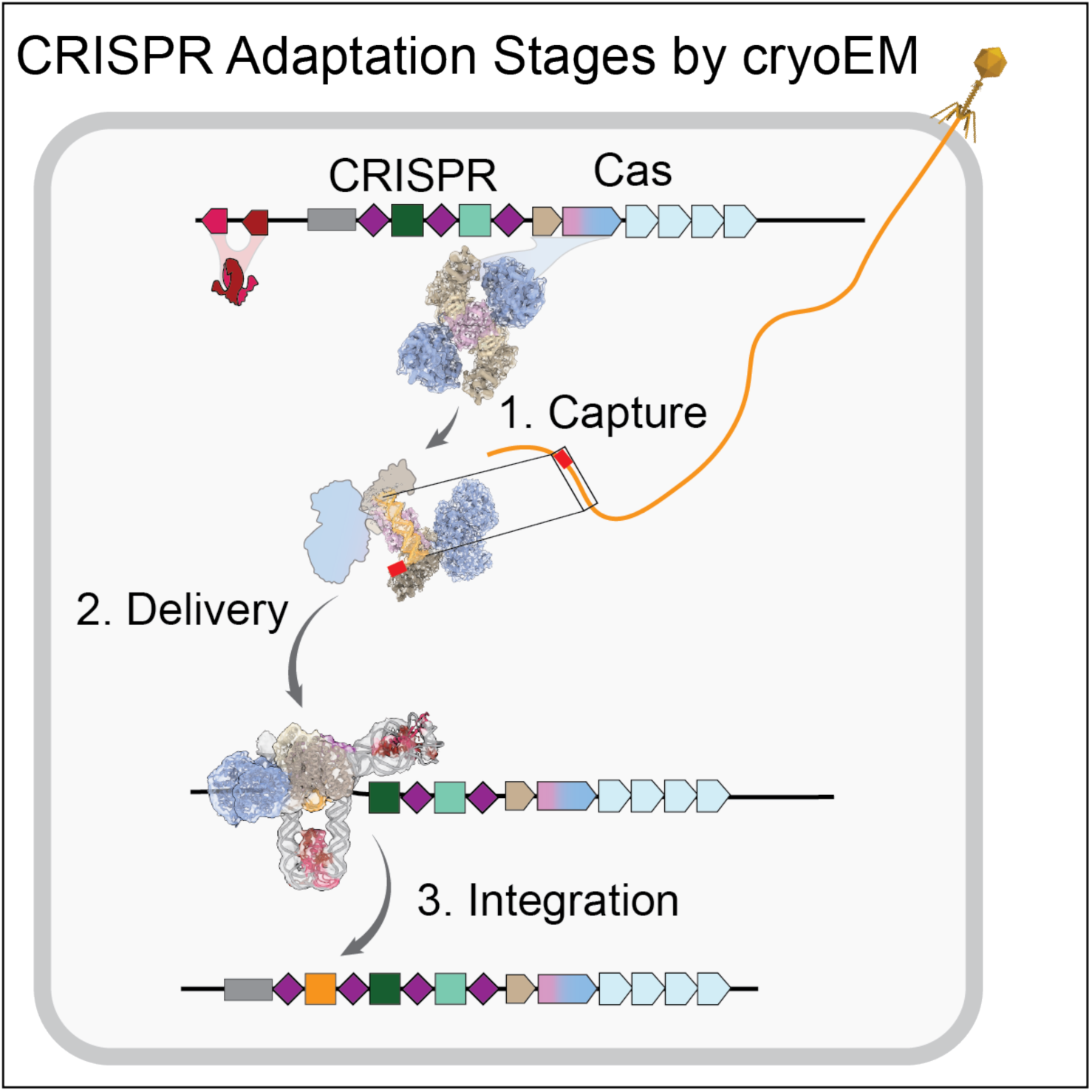

**HIGHLIGHTS:**

- A positively charged channel on the Cas1-2/3 complex captures fragments of DNA
- A loop in the RecA1 domain controls access to the Cas3 nuclease active site
- Foreign DNA binding allosterically regulates access to additional DNA binding sites
- Distortion of the CRISPR repeat sequence licenses complete foreign DNA integration

## INTRODUCTION

Adaptive immunity requires genomic recombination to generate molecular memories of pathogens. In both V(D)J recombination in eukaryotes and CRISPR-Cas adaptive immunity in prokaryotes, viral-derived transposases have been independently domesticated to fight foreign invaders^1–6^. However, the mechanism of adaptive immunity differs significantly. In jawed vertebrates, the RAG recombinase splices together fragments (i.e., V,D and J) of genomic DNA, but these adaptive rearrangements are not heritable^1,7–9^. In contrast, prokaryotic Cas integrases capture and insert short fragments of foreign DNA into CRISPR loci, creating an updated and heritable record of circulating viral predators^10–13^.

Cas1 is a striking example of a selfish transposase gene repurposed for host defense^3,3,14^. CRISPR-Cas systems, nearly all of which contain Cas1, are found in ∼90% of archaea and 40% of bacteria^3,14–16^. Cas1 no longer mobilizes a selfish genetic element but protects the cellular host by providing the CRISPR interference machinery with updated guide sequences derived from selfish genetic elements^17–21^. During the adaptation stage of CRISPR-Cas adaptive immunity, Cas1 and Cas2 assemble into a heterohexameric complex that captures and integrates fragments of foreign DNA into the CRISPR array^6,10,17,18,21–26^. These fragments often contain a 2-5 bp protospacer adjacent motif (PAM) that functions to distinguish self from non-self DNA and facilitates directional integration into the CRISPR array^27^. In some CRISPR systems, dedicated Cas proteins recognize and trim the PAM prior to integration, while in other systems this function is performed by non-Cas host proteins such as DnaQ exonucleases^28–33^. However, the mechanism of PAM trimming remains unknown in many CRISPR systems that lack the canonical PAM-trimming protein Cas4^22,32,33^.

Cas3 is a nuclease/helicase found in all type I CRISPR systems^16,34^. The nuclease/helicase generates foreign DNA fragments by degrading foreign DNA identified by the CRISPR RNA-guided surveillance complex (i.e., Cascade or Csy complex)^23,35–40^. Cas3 functionally couples the interference and adaptation stages of CRISPR immunity by generating dsDNA fragments enriched in PAMs^22,34,41,42^, which serve as substrates for integration^20,41,42^. This link between generating (Cas3) and integrating (Cas1 and 2) fragments of foreign DNA is further highlighted in type I-F CRISPR-Cas systems, where Cas3 is genetically fused to the C-terminus of Cas2^16^. Cas1 and Cas2/3 assemble into a two-fold symmetric propeller-shaped heterohexamer where Cas1 represses the Cas2/3-nuclease activity through electrostatic interactions at a single interface^20,41^. However, the Cas1-2/3 complex is anticipated to adopt distinct conformational states that support its diverse roles in target degradation, foreign DNA capture, site specific DNA delivery, and foreign DNA integration^20,41,43^.

Here we determine cryo-EM structures of the type I-F Cas1-2/3 integration complex from *Pseudomonas aeruginosa* in multiple conformational states. These structures explain how Cas1-2/3 recruits short fragments of foreign DNA to a positively charged channel on one face of the complex. Foreign DNA binding to this face functions as an allosteric switch that triggers conformational rearrangements, exposing DNA binding sites previously occluded by Cas1 and Cas3. Cas3 sterically regulates the integrase but does not trim the PAM. Nonetheless, PAM trimming is an essential prerequisite for positioning foreign DNA in the Cas1 active site and completion of the second transesterification reaction. Taken together, these cryo-EM structures illustrate how the Cas1-2/3 complex orchestrates foreign DNA capture, site specific delivery, and integration of new DNA into the bacterial genome.

## RESULTS

### Cas1-2/3 forms a DNA capture complex

To determine how the Cas1-2/3 integrase captures fragments of foreign DNA, we set out to determine the first high-resolution structure of a Cas1-2/3 complex in the absence of nucleic acid. Cas1 and Cas2/3 proteins from the *PA14* strain of *Pseudomonas aeruginosa* were co-expressed in *Escherichia coli* BL21 cells. An amino-terminal Strep tag on Cas1 pulls down untagged Cas2/3 in a stable complex that elutes from a size exclusion column as a single peak with an estimated molecular weight of ∼380 kDa, corresponding to a heterohexameric assembly with four Cas1 subunits and two Cas2/3 subunits (**Figure 1A** and **S1**)^20,41^.

**Figure 1.**
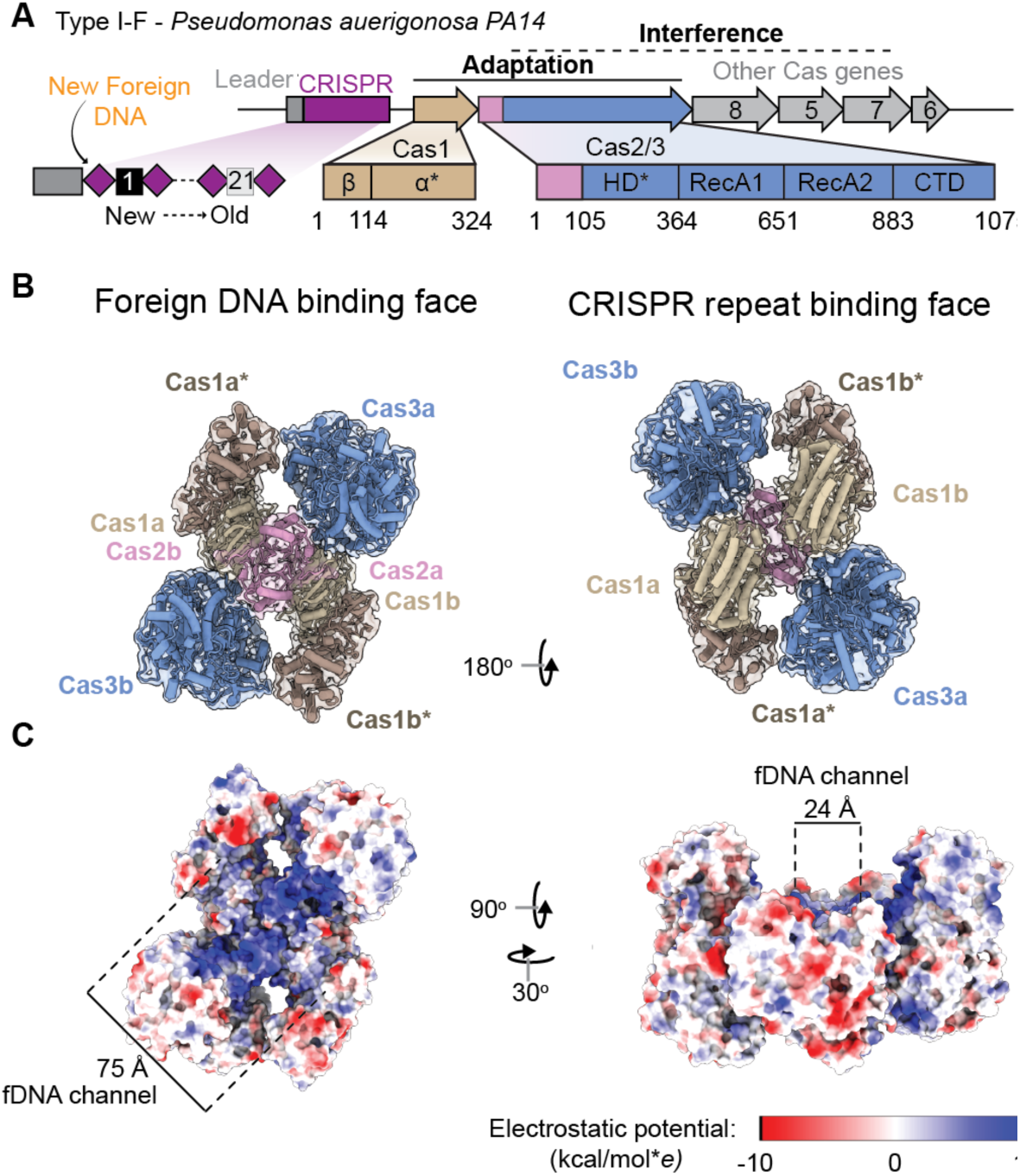
One side of the Cas1-2/3 integrase has a positively charged foreign DNA-binding channel. (A) Schematic of the CRISPR locus (purple) and *cas* genes in the type I-F system from *Pseudomonas aeruginosa.* The asterisks (*) indicates nuclease domains (B) A 3.3 Å resolution cryo-EM structure of the Cas1-2/3 heterohexamer reveals a propeller shaped complex with C2 symmetry. One face of the complex is involved in binding foreign DNA, while the other binds the CRISPR repeat. The asterisks (*) indicates the Cas1 subunits where transesterification will occur. Density map displayed at 80% transparency (threshold=0.109), with the atomic model docked inside and colored by subunit. (C) A large patch of basic residues is present on one face of the complex. This patch forms a channel 24 Å wide and 75 Å long. Atomic model is colored by electrostatic potential using the ChimeraX command “coulombic” (kcal/mol\**e* at 298 K).

Purified Cas1-2/3 was concentrated to 12 μM, spotted on cryo-EM grids, blotted, and plunge frozen in liquid ethane. Frozen grids were imaged using a 300 keV cryo-electron microscope equipped with an energy filter. Grid squares with thin ice and a dense particle distribution were used to collect 3,944 movies. After preliminary motion correction, CTF-correction, blob picking, and 2D classification, *ab inito* reconstruction and non-uniform refinement were used to reconstruct an initial volume that was used as a template for particle picking. 303,250 template-picked particles were extracted from 3,293 cryo-EM movies to determine a 3.3 Å resolution structure of the complex (**Figure 1B-C** and **S1)**. To build an atomic model of the complex, an AlphaFold3-predicted structure of the complex was fit into the electron density map using ChimeraX, followed by iterative refinement in Isolde, Phenix, and Coot^44–46^. This approach resulted in an atomic model across 96% of Cas1-2/3 residues with 3,138 unambiguously positioned sidechains (**Table S1**).

Cas1-2/3 is shaped like a four-bladed propeller. The Cas1 homodimers and Cas3 domains form the blades around the central Cas2 homodimer (**Figure 1B**). In type I-F systems, the small Cas2 subunits (∼12 kDa) are uniquely fused to the large Cas3 protein (∼109 kDa) by a linker sequence (residues 95-104), which is not resolved in the density due to its flexibility. To understand if the fusion of the large Cas3 effector to the C-terminus of Cas2 alters the core integrase architecture compared to previously described assemblies of Cas1-2, we compared the Cas1-2/3 structure to previously determined structures (**Figure S2** and **S3**)^26,47^. All Cas2 proteins described to date have the same N-terminal ferredoxin-like fold (βαββαβ topology) and a fifth anti-parallel beta strand that completes the homodimer interface^26,47–49^. This fifth beta strand is provided by the same molecule in type I-E systems and by the opposing Cas2 molecule in other CRISPR systems^47,50^. Whether the fifth beta strand is provided in *cis* or *trans* corresponds to two points of contact with Cas1 in type I-E systems (*cis*) or one Cas1 interaction in type I-A systems (*trans*) (**Figure S2** and **Table S2)**^51,52^. Like the I-E system, the I-F Cas2 provides the fifth anti-parallel beta strand in *cis* and interacts with two Cas1 molecules to create a molecular ruler that measures DNA sequences of similar lengths in both type I-E and type I-F systems^19,26^. The C-terminal fusion of Cas3 to Cas2 does not fundamentally alter the core Cas1- Cas2 integrase architecture (**Figure S2, S3)**.

Previous work established that four residues in each Cas2/3 molecule coordinate the phosphate backbone of double-stranded B-form foreign DNA during integration^43^. These four Cas2 residues (K11, R18, T93, and C94) form the edges of a 24 Å-wide positively-charged channel, which we designate the “foreign DNA binding face” (**Figure 1B-C** and **S4**). Histidine residues in Cas1 (H25) are like wedges on either end of the channel (**Figure 1C** and **S4**). These histidines split dsDNA and direct 3’ ssDNA ends down the electrostatic funnel toward the Cas1 active site^43^. The length (75 Å) and the width (24 Å) of the “foreign DNA binding channel” can accommodate 22 base-pairs of B-form DNA and is the only face of the Cas2 homodimer accessible for DNA binding (**Figure 1**). The opposing “CRISPR repeat binding face” of Cas2 is obscured by Cas1 homodimers (**Figure 1B**). Previous biochemical analysis reveals that a K11D charge-swap mutation in Cas2 prevents integration of foreign DNA into the CRISPR array^43^.

Thus, the cryo-EM structure of Cas1-2/3 in the absence of DNA reveals that the solvent accessible face of Cas2 forms a positively charged foreign DNA binding channel.

To understand if the residues involved in binding and guiding DNA to the Cas1 active sites are conserved, we calculated conservation scores for type I-F Cas1 and Cas2/3 proteins from NCBI’s non-redundant database of complete bacterial and archaeal genomes (see Materials & Methods). This analysis revealed a striking pattern of conservation across the foreign DNA binding face of Cas1-2/3 and low conservation on other solvent-exposed surfaces (**Figure S4**). Three of the four DNA binding residues on Cas2 (K11, R18, and T93) are conserved, as well as the positively charged residues in the electrostatic funnel that leads to the Cas1 active site (R72, R76, R293, and R259) (**Figure S4**). To understand if the propeller assembly itself is a conserved feature of the Cas1-2/3 complex, we used AlphaFold3 to predict the structure of four distinct Cas1-2/3 complexes. Each of these predicted structures were aligned to the experimentally determined structure (**Figure S5**). All predicted structures of the Cas1-2/3 complex preserved the four-bladed propeller architecture observed in the experimentally determine structure (**Figure S5)**. These comparisons predict that both the Cas1-2/3 foreign DNA binding channel and propeller assembly are both conserved structural features of I-F CRISPR systems.

### Cas3 regulation by a conserved gate and latch

Cas3 is an HD nuclease and a superfamily 2 (SF2) helicase that degrades targeted foreign DNA during type I CRISPR interference^34,40^. Previous studies show that Cas1 inhibits the nuclease activity of Cas3, while Cas1-2/3 recruitment to target-bound Cascade relieves Cas3 inhibition and leads to rapid DNA degradation^35,36,41,53^. The cryo-EM structure of Cas1-2/3 reveals the RecA1 and RecA2 helicase domains stack to form a solvent-accessible cleft for ATP binding and hydrolysis, while the HD nuclease domain associates with the RecA1 domain, as in previously determined structures of Cas3^35–37,54^. (**Figure 2A-B)**. In the absence of Cascade, a loop from RecA1 (residues 483-511) covers the HD active site like a closed gate (**Figure 2B**). At the top of this loop, R148 and K127 of the HD domain form hydrogen bonds with G498 and E500 (**Figure 2C**). These hydrogen bonds hold the gate over weak unmodelled density, which we attribute to the heterogenous occupation of two divalent metal cations in the HD active site that are known to be critical for HD nuclease function (**Figure 2C**)^35,40,55^. Examining the alignment of Cas2/3 fusion sequences (n=685) revealed a well-conserved motif (GSES, residues 498-501) that appears to function as a latch (**Figure 2D**). Notably, when Cas2/3 associates with Cascade, the single-stranded R-loop displaces the latch and the gate flips open, creating a direct path to the HD active site where ssDNA cleavage occurs (**Figure 2E**)^56^. The RecA1 gate hinges at a conserved motif found across SF2 helicases (motif 1b), including Cas3s, suggesting that this hinge may be functionally conserved in SF2 helicases^34^. Auto- inhibition of the HD nuclease of Cas3 by insertions in the RecA1 helicase has been described in the type I-A system of *Pyrococcus furioisus*^57^, and a similar gate-like feature is displaced upon ssDNA binding in crystal structures of Cas3 from type I-E systems^35,58^. While the gate-like insertion in RecA1 is structurally conserved, the amino acid sequence differs across systems.

**Figure 2.**
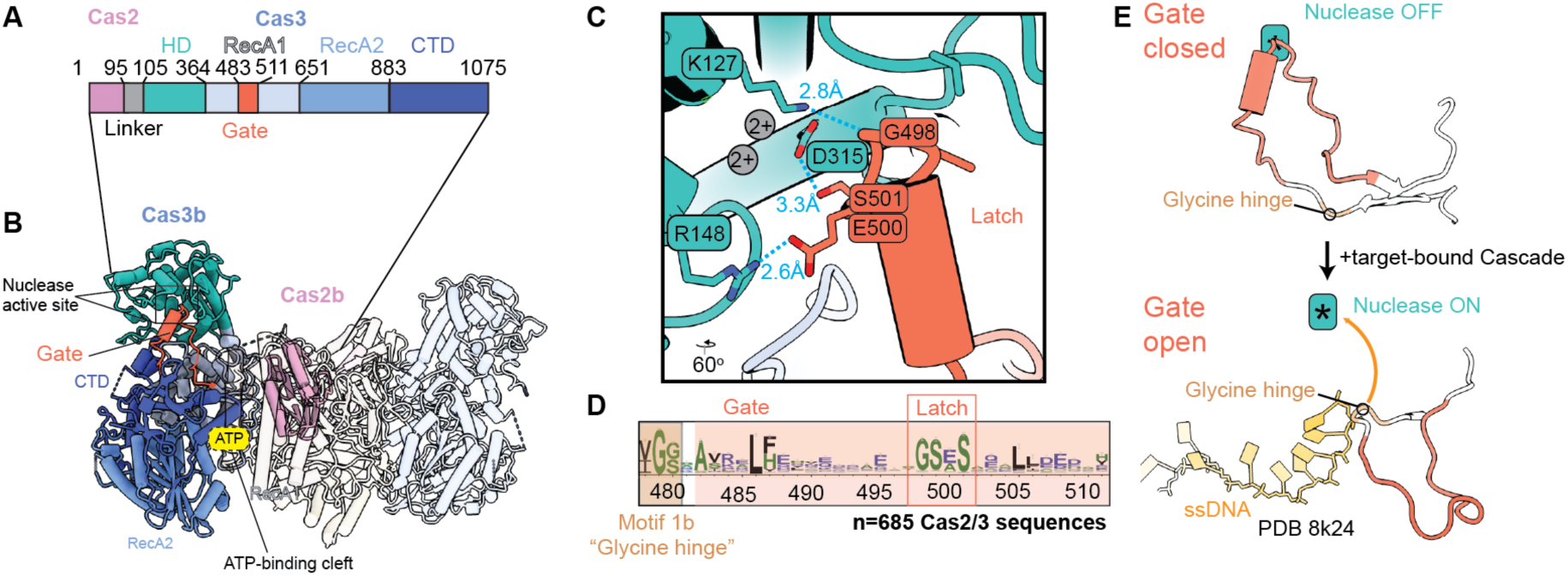
A conserved gate-like feature obscures access to the Cas3 HD nuclease active site. (A) Domain architecture of the Cas2/3 fusion. (B) A RecA1 insertion (residues 483-511, colored orange) protrudes into the HD-nuclease active site of Cas3. One Cas2/3 molecule colored by domain, while all other molecules are displayed at 90% transparency. The yellow ATP label indicates the ATP binding site between RecA1 and RecA2. (C) Residues of the HD domain (K127, R148, D315) interact with the gate residues (G498, E500, S501) latching the gate over HD active site, where two divalent cations are positioned in the active site (grey circles, labelled 2+). Hydrogen bonds are shown in turquoise, with the distance between hydrogen-bonded atoms indicated. (D) The RecA1 gate and latch contains two sequence motifs, motif 1b at the glycine hinge (G479) and the latch motif (GSES, residues 498-501), that are conserved in type I-F systems. WebLogo built from a sequence alignment of n=685 Cas2/3 sequences. (E) The R-loop formed by Cascade during RNA-guided strand invasion displaces the HD gate during interference (PDB 8k24) reveals the HD gate is flipped out of the active site, rotating from the glycine hinge (motif 1b). Single-stranded DNA (ssDNA) travels along the path previously occluded by the gate to the HD active site, indicated with an asterisk (*).

Collectively, this work and prior observations suggest that Cas3 RecA1 insertion sequences gate access to the HD nuclease, which may help prevent off-target DNA degradation by Cas3.

### Cas3 blocks leader binding sites of Cas2

Previous work has shown that Cas2 is essential for bending the CRISPR leader during foreign DNA integration in type I-F systems and that residues R55 and N56 are critical for this interaction^43^. During integration, R55 intercalates between deoxyribose sugars and N56 stabilizes the phosphate backbone at the conserved type I-F inverted repeat motifs^19,43^. Based on these interactions, we named the face of Cas2 containing these residues the “leader binding site”. In the absence of DNA, the leader binding site is masked by a network of hydrogen bonds and salt bridges between residues in Cas2 (R54, R55, K50 and E71), and the RecA1 domain of Cas3 (N373, D472, D473, T517, E519, D520, R525, R529) (**Figure 1, S3** and **S6**). Residues R55 and R54 of Cas2 form the most extensive network of hydrogen bonds with the RecA1 domain (**Figure S6**). Both the cryo-EM map and AlphaFold predictions reveal the position of Cas3 over the leader recognition sites is further stabilized through an elaborate network of hydrogen bonds with Cas1 dimers on opposite ends of the complex (**Figure S6**). Collectively, these interactions stabilize Cas3 over the leader binding sites on Cas2.

### Foreign DNA capture exposes DNA binding sites

The propeller-shaped conformation of Cas1-2/3 alone is drastically different from the conformation of Cas1-2/3 bound to a synthetic integration intermediate (**Figure 1**)^43^. To determine how the Cas1-2/3 complex positions short DNA fragments for PAM trimming and integration, we incubated Cas1-2/3 with a 32 bp fragment of foreign DNA. This DNA fragment contained 22 bp of dsDNA and 5 nts of splayed single-stranded ends with two additional 3’ GG PAM nucleotides added to one strand (**Figure S7**). The DNA-bound Cas1-2/3 complex eluted from a size exclusion column with an estimated molecular weight of 415 kDa, which is consistent with a 1:1 stoichiometry of Cas1-2/3 and foreign DNA (**Figure S7)**. A cryo-EM dataset of 5,552 movies was collected and processed to determine a 3.31 Å reconstruction of this DNA-bound Cas1-2/3 complex (**Figure 3, S7** and **Table S3)**. Conformational heterogeneity and asymmetry were notable features of this dataset compared to data collected on the Cas1- 2/3 complex without DNA. However, during processing, no classes corresponding to the DNA-free complex were observed, indicating that foreign DNA stably associates with Cas1-2/3. The primary source of heterogeneity was the position of Cas3 relative to Cas1-2 (**Figure S7)**. This heterogeneity was not a result of particle picking bias, since blob picking followed by cryoSPARC template picking, crYOLO, Topaz, or cryoSPARC’s DeepPicker all yielded particle stacks of similar sizes and heterogeneity^59–61^. An AlphaFold3 prediction of the DNA-bound Cas1-2/3 complex positioned Cas3 in two different orientations relative to Cas1-2, providing further evidence that DNA binding to the foreign DNA binding face in fact destabilizes Cas3 positioning (**Figure S8**).

**Figure 3.**
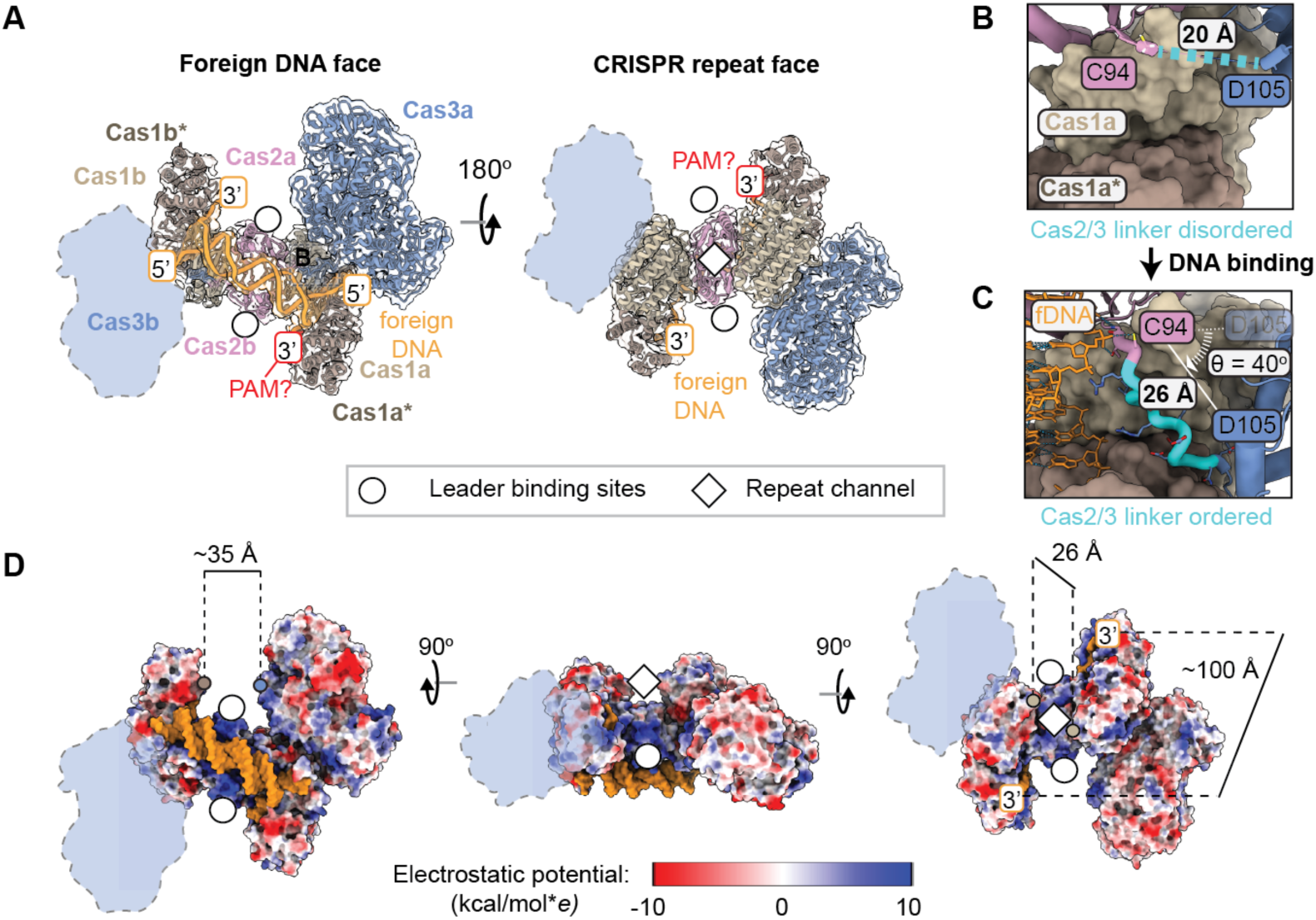
Foreign DNA triggers a conformational rearrangement that exposes additional DNA binding sites. (A) Atomic model colored by subunit docked into ∼3.3 Å density map from a non-uniform refinement shown at 80% transparency (threshold=0.178). One Cas3 (shown as an outline) is conformationally flexible. (B) The linker between C94 and D105 in Cas2/3 is flexible and unresolved in the absence of foreign DNA (shown as a dashed line in cyan). (C) Foreign DNA binding lengthens and orders the 10 amino acid linker between Cas2 (first residue: C94) and Cas3 (last residue: D105), changing the position of Cas3 relative to Cas1-2 (D) Foreign DNA binding exposes two leader binding sites and a basic channel for the CRISPR repeat. The leader binding sites sit at the base of a 35 Å channel, measured from D388 on Cas3a (blue dot) to G236 on Cas1b* (brown dot). The length of the repeat channel was measured from Cas1a* H271 to Cas1b* H271 (∼100 Å, positioned beneath the 3’ label). The width of the CRISPR repeat binding channel was measured from R169 Cas1a to R169 on Cas1b (beige dots) . Atomic model is colored by electrostatic potential (kcal/mol\**e* at 298 K).

To build an atomic model of the complex, the AlphaFold3 predictions of the Cas3 domain and the Cas1-2 hexamer bound to dsDNA were independently rigid-body fit into the density in ChimeraX followed by jiggle fitting in Coot^45^. This model was used as a starting point for iterative refinement in Isolde, Phenix, and Coot^44–46^, leading to a map with 66% of main chain residues unambiguously modeled (**Table S3**). This model captures both Cas1 homodimers, the Cas2 homodimer, and foreign DNA, but only one of the two Cas3 lobes (**Figure 3A**). Given that Cas2 and Cas3 are fused, the presence of Cas2 and absence of Cas3 in the cryo-EM density map indicates that Cas3 is averaged out during processing because it exists in multiple distinct conformations. Thus, this structure reveals that DNA binding triggers release of Cas3 from the leader binding sites of Cas2. One Cas3 domain is stabilized in a new position against the Cas1 dimer while the opposing Cas3 rotates freely.

The DNA fragment preferentially binds to the foreign DNA binding face of Cas1-2/3 (**Figure 3**). DNA binding triggers conformational changes that expose three additional DNA binding sites involved in integration (**Figure 3**). The individual domain architectures of Cas2 and Cas3 remain largely unchanged after binding foreign DNA (RMSD < 1 Å across 94 unpruned Cα pairs for the Cas2 domain, 972 unpruned Cα pairs for the Cas3 domain), but the positioning between domains changes dramatically. Upon foreign DNA binding, the linker between the Cas2 and Cas3 domain (residues 95-104) rigidifies, extends, and rotates to reposition Cas3 (**Figure 3B-C**). This rearrangement breaks all the previous Cas3 interactions with both Cas1 homodimers and Cas2, and establishes new interactions between Cas3 and a single Cas1 homodimer (**Figure 1 3,** and **S6**). The new Cas3 position creates a 35 Å wide positively charged channel between Cas3 and Cas1 that exposes the CRISPR leader binding sites on both sides of the Cas2 homodimer (**Figure 3D**, indicated with white-filled circles). The Cas1 dimers each rotate out and down, which opens a 26 Å wide positively charged channel that we designate the “CRISPR repeat channel” based on previous work (**Figure 3D**, white diamonds)^17,33,43^. The Cas2 homodimer forms the floor of this channel. The transesterification sites of the active Cas1 subunits are 100 Å apart at either end of the CRISPR repeat channel, while the inactive Cas1 subunits form the walls of the channel (**Figure 3**). The domain arrangements of Cas1-2/3 bound to foreign DNA are nearly identical to the domain arrangements of the previously determined type I-F integration complex^43^, revealing that foreign DNA binding is sufficient to position the Cas1-2/3 complex for integration.

### DNA positioned for PAM trimming and integration

Despite the presence of three additional exposed positively charged DNA binding sites, nucleic acid density is only observed on the foreign DNA binding face (**Figure 3** and **S7**). The absence of nucleic acid density at the other DNA binding sites suggests that recruitment of nucleic acid to the leader binding sites and repeat channel requires sequence- or structure-specific interactions to coordinate assembly of the integration complex at the CRISPR locus (**Figure 3** and **S7)**. As in the previously determined structure of the integration complex^43^, 22 base pairs of the double-stranded DNA is positioned in the foreign DNA binding channel by residues K11 and R18 of Cas2 through hydrogen bonds with the phosphate backbone (**Figure S9**). The H25 wedge splits the double stranded nucleic acid and routes the 3’ strand down a positively-charged channel leading to the Cas1 active site (**Figure S9**), where the 3’ ends of foreign DNA are positioned for transesterification. This positioning is further stabilized by hydrogen bonds between the backbone nitrogens of residues T93 and C94 of Cas2/3 with the phosphate backbone of the foreign DNA (**Figure S9**). The residues involved in positioning the foreign DNA are conserved in type I-F systems, highlighting their functional importance (**Figure S9**).

The foreign DNA substrate contained an additional 3’ GG dinucleotide PAM on one end (**Figure S7**). The PAM is critical for directional insertion of foreign DNA into the CRISPR array, but the protospacer-adjacent motif must be removed prior to or during DNA integration^29,30,33,43^. However, the mechanism of PAM trimming is unknown in type I-F CRISPR systems. We hypothesized that Cas3 might play a role in PAM trimming because it contains a nucleolytic HD domain and undergoes substantial rearrangement relative to Cas2 and Cas1 upon foreign DNA binding. However, after foreign DNA binding to the Cas1-2/3 complex, the Cas1 transesterification sites are positioned ∼80 and 100 Å away from the HD domain (**Figure 3 and S9**). Though Cas3 is both an endonuclease capable of nicking ssDNA and a 3’ to 5’ exonuclease^40^, it’s positioning in the Cas1-2/3 complex precludes a role for it in PAM trimming without additional conformational changes.

DNA-induced structural rearrangements expose the faces of Cas1 and Cas2 that have been shown to interact with PAM-trimming exonucleases in other CRISPR systems (**Figure S9**)^31,33^. To understand if Cas3 would sterically block access to the PAM, we docked a previously determined structure of Cas1-Cas2-Cas4 bound to foreign DNA onto the atomic model of the Cas1-2/3 complex bound to foreign DNA (**Figure S9**). While the I-F system doesn’t contain a Cas4 protein, it’s role in PAM trimming serves as a structural proxy for the unidentified nuclease responsible for PAM trimming in type I-F systems. Docking reveals that the 35 Å channel between Cas1 and Cas3 accommodates Cas4 without steric clashes between Cas3 and Cas4 (**Figure S9**), demonstrating that foreign DNA-induced conformational changes may also facilitate the recruitment of an unidentified PAM trimming nuclease by exposing a conserved face of Cas1 that positions the 3’ PAM for trimming (**Figure S9**). Weak density in a global refinement of the foreign DNA-bound Cas1-2/3 complex refined to 3.3 Å resolution suggests that the PAM is positioned in the Cas3-stabilized side of the complex. To unambiguously assign PAM nucleotide positions, each Cas1 dimer was refined separately using local masks.

Surprisingly, the two Cas1 dimer volumes did not differ significantly. This result suggests that the nucleic acid provided did not completely lock the PAM in a specific orientation at either Cas1 active site. However, the DNA-induced asymmetry suggests a role for the PAM in asymmetric stabilization of the complex.

### The PAM alters integration states

The Cas1-2/3 integrase is a C2 symmetric heterohexamer (**Figure 1**)^17,18,30,43^. This symmetric integrase must home in on the CRISPR leader in an orientation that ensures that the spacer is integrated into the CRISPR array in a direction that results in transcription of a CRISPR RNA that is complementary to an invading DNA sequence adjacent to the PAM^17,43^. Integration Host Factor (IHF) is also critical for introducing folds in the CRISPR leader recognized by Cas1-2/3^43^. Previous *in vitro* biochemical experiments performed using purified components of the type I-F system revealed that the PAM stalls integration after the first transesterification reaction^17,43^. To understand if the PAM functions to orient the integrase onto the CRISPR array, we determined the structures of Cas1-2/3 integrating foreign DNA with and without a PAM into the CRISPR array (**Figure 4** and **S10-11**). DNA substrates with or without a PAM were incubated with purified Cas1-2/3 and then mixed with purified IHF and a 200 bp fragment of DNA containing the 5’ CRISPR leader immediately upstream of a repeat-spacer sequence. This mixture was incubated at 35 °C for 25 minutes then subjected to size exclusion chromatography. The purified integration complex was concentrated, spotted on grids, blotted, and plunge frozen in liquid ethane. Two datasets of 6,677 movies (PAM-containing) and 5,462 movies (no PAM) were collected, and the previously determined integration complex (PDB 8FLJ) was used as a template for particle picking^43^. This template picking strategy identified 646,531 unique particles in the PAM-containing dataset and 615,271 unique particles in the no PAM dataset. From these particles, five distinct mid-resolution electron density maps of the integration complex were refined: two maps from the PAM-containing dataset (3.9 Å and 4 Å), and three from the no PAM dataset (3.9 Å, 3.9 Å and 3.8 Å) (**Figure 4, S10-11** and **Table S3-S4**).

**Figure 4.**
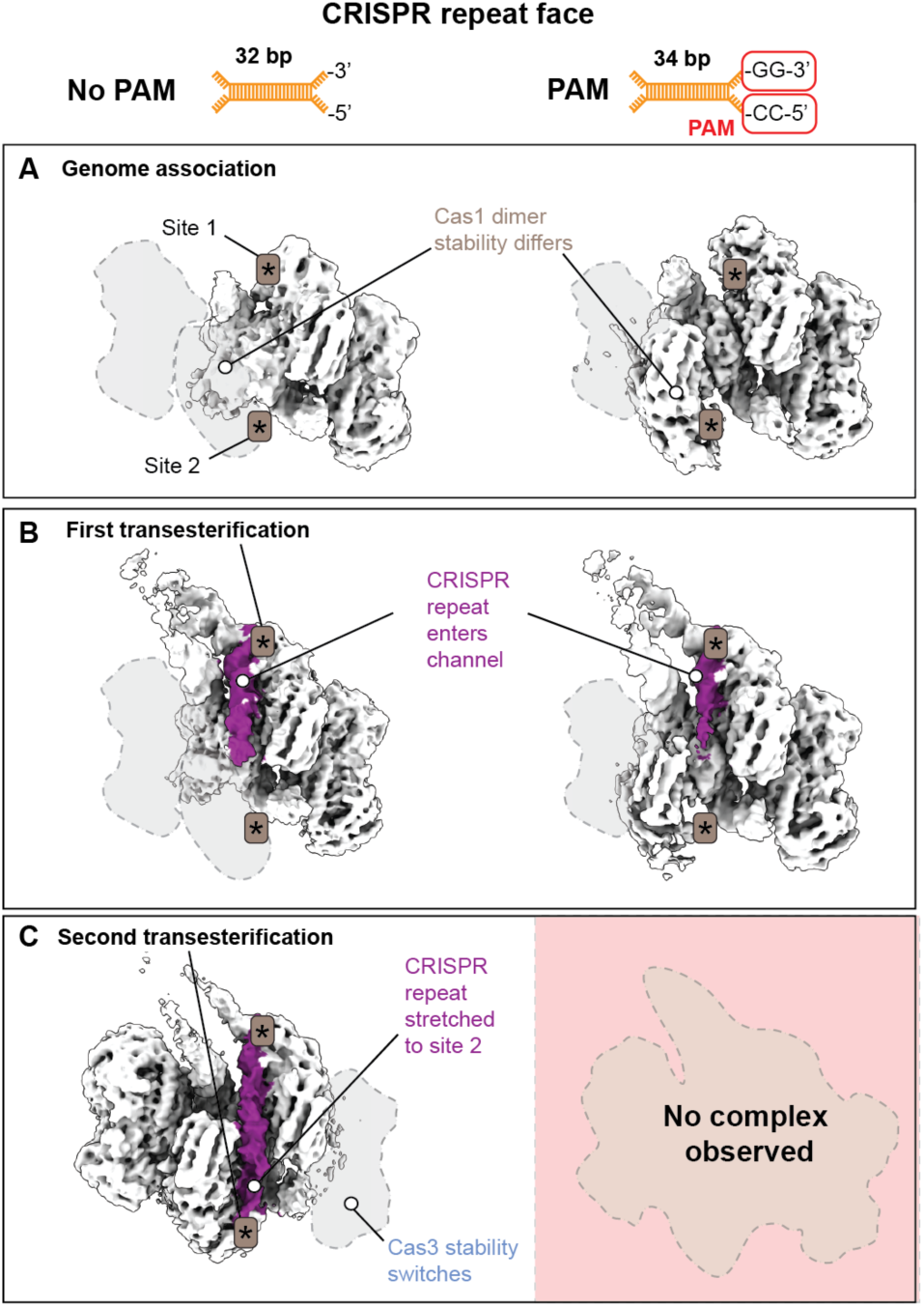
Foreign DNA length affects complex stability and sequential transesterification. (A-C) Cryo-EM density maps of the Cas1-2/3 complex integrating foreign DNA fragments into the CRISPR repeat with or without a PAM (threshold=0.103). Density for the CRISPR repeat is colored purple. Asterisks (*) indicate the Cas1 transesterification sites. Dashed outlines indicate subunits missing due to conformational flexibility. (A) Density maps from non-uniform refinements of in the presence (82,751 particles) and absence (n=121,485 particles) of a PAM captured in a genome-associated, pre-transesterification state. (B) Density maps from non-uniform refinements in the presence (n=94,204 particles) and absence (n=97,437 particles) of PAM after the first transesterification reaction covalently links the CRISPR repeat to the foreign DNA fragment (C) In the absence of a PAM, density corresponding to the CRISPR repeat is positioned in the active site of the Cas1 subunit responsible for the second transesterification (n=93,957 particles). The repeat stretches across the channel.

The presence of the PAM in the foreign DNA fragment did not change how Cas1-2/3 initially associates with the CRISPR leader (**Figure 4B-C**). The integrase folds the CRISPR leader and the leader-repeat junction is positioned into the first transesterification site (Site 1) in both the presence and absence of PAM (**Figure 4D-E**). In all volumes, the short, splayed-end fragments of DNA bind to the foreign DNA binding face of Cas1-2/3 and there is no evidence for the longer DNA fragments bound at this location, providing structural confirmation that Cas1-2/3 has a length preference on the foreign DNA binding face^20^. As observed in the delivery complex, the integration complex is asymmetrically stabilized, with density for one Cas3 averaged out due to conformational flexibility in all volumes (**Figure 3,4**, **S10, and S11**).

The presence of the PAM introduced two notable differences in the integration complex. First, the second transesterification reaction site (Site 2) is resolved in the presence of a PAM due to stabilization of the corresponding Cas1 dimer (**Figure 4A**). Second, when a PAM is present, the repeat fails to extend all the way through the repeat channel (**Figure 4B-C**). In the absence of a PAM, the repeat stretches through the repeat channel to the second transesterification site. When the repeat channel is fully occupied, the Cas1 dimer housing the second transesterification site appears stabilized, along with the previously unresolved Cas3b domain. On the opposite side of the complex, the previously stabilized Cas3 is averaged out, presumably due to conformational heterogeneity (**Figure 4C**). These structures explain previous biochemical data showing that the presence of a PAM blocks the second transesterification reaction but not the first^43^. The PAM does not alter initial integration complex assembly but blocks the formation of structural states required for the second transesterification reaction to occur.

### Full integration distorts the repeat

In order for the second transesterification reaction to occur, the Cas1-2/3 complex must dock the first Cas1 transesterification site at the leader-repeat junction (Site 1) and the second Cas1 transesterification site (Site 2) at the spacer-repeat junction^19,43^. To ensure complete duplication of the repeat on either side of the newly integrated foreign DNA, both the leader-repeat junction and the spacer-repeat junction must be correctly positioned in the Cas1 transesterification sites. To understand if the repeat sequence forms secondary structures that facilitate this positioning, AlphaFold3 was used to predict the structure of the repeat sequence alone. The 32 bp repeat sequence adopts linear B-form conformation that is ∼7 Å too short to stretch from Site 1 to Site 2, revealing that the repeat must be distorted during sequential transesterification (**Figure 5A**). To understand how the repeat is distorted, the linear B-form Alphafold3 prediction was docked into the repeat density from the only class containing density along the full repeat channel (**Figure 4C** and **5A**).Though the conformational heterogeneity induced by repeat binding in the channel precludes construction of an atomic model of this state (**Figure 5B**), a local refinement of the low-resolution volume compared to the predicted nucleic acid structure highlights several important features of the CRISPR repeat during integration. Leaving the first transesterification site, the repeat adopts a pseudo-B-form helical conformation for ∼25 Å (∼6-7 bp) (**Figure 5B-D**). A kink in the density marks the beginning of a distorted region in which the helical pitch of B-form DNA is bent into the narrow central region of the channel (**Figure 5B-D**). A second kink marks a transition back to pseudo-B-form DNA as the repeat approaches Site 2 (**Figure 5C-D**). These two kinks change the trajectory of the CRISPR repeat in the repeat channel, bending the DNA ∼28° over the top of the Cas2 dimer (**Figure 5B)**. The repeat distortion also introduces a ∼3° shift towards the second transesterification site **(Figure 5B**). Notably, positively charged residues that line the CRISPR repeat channel are conserved in type I-F systems, highlighting the importance of residues in the CRISPR repeat channel positioned to distort the repeat (**Figure S12**).

**Figure 5.**
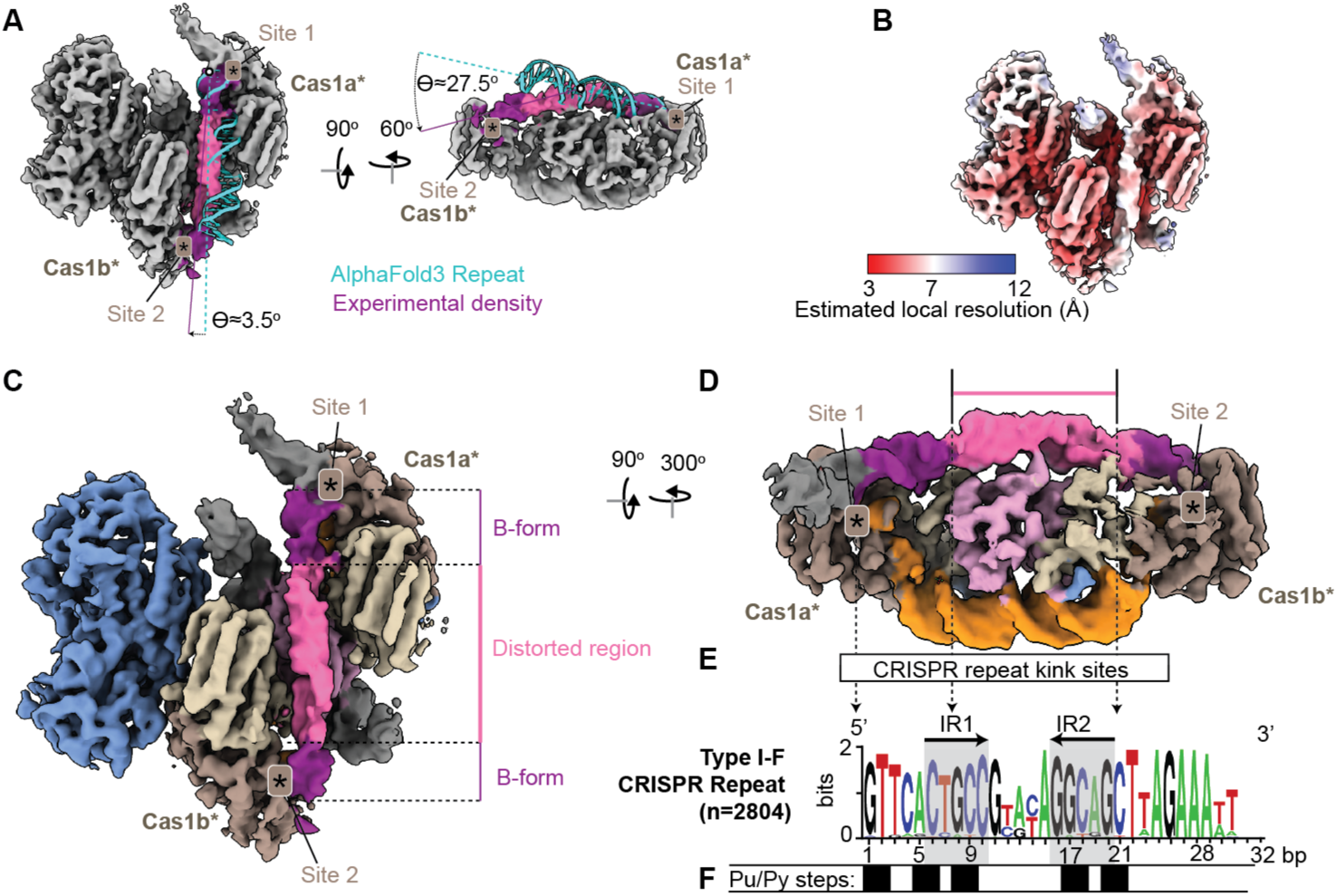
The repeat channel distorts the CRISPR repeat. (A) An AlphaFold3 prediction of the 28 bp repeat (cyan) from the PA14 strain of *P. aeruginosa* was rigid body fit into the B-form section of the density map projecting from Site 1 (colored purple). The experimental density for the repeat is twisted 3.5° off-center and bent ∼27.5° down into the second transesterification site (Site 2) compared to the AlphaFold3 prediction. Cas1 active sites where transesterification occurs are indicated with an asterisk (*), and the density is displayed at threshold=0.171. B-form regions of repeat density are colored purple, and distorted regions are colored pink. (B) Local resolution estimation of the repeat-containing volume (n = 27,573 particles) displayed at threshold 0.171 (C) The repeat is distorted in the middle of the repeat channel as it extends from the first transesterification site (Site 1) to the second site (Site 2). The distorted region is displayed in pink, while the B-form region of the repeat is displayed in purple. Cas1 active sites are indicated with an asterisk (*), and the density is displayed at threshold=0.171 and colored by protein subunit. (D) Side view of the distorted repeat (threshold=0.171, protein subunits that obscure the repeat at this view are erased for clarity). Two kinks in the density flank the distorted region, and approximately align with conserved purine-pyrimidine steps (indicated by dashed arrows pointing to panel E). Cas1 active sites are indicated with an asterisk (*). (E) A logo plot showing the conservation of the type I-F CRISPR repeat across 2804 aligned type I-F repeat sequences (28-32 bp). Letter height corresponds to bit-score from 0 (not conserved) to 2 (conserved). Inverted repeats are indicated with grey boxes and black arrows indicate direction of inversion. (F) There are five conserved di-nucleotide purine (Pu) to pyrimidine (Py) steps in the type I-F CRISPR repeat at positions 1-2, 5-6, 8-9, 17-18, and 20-21, indicated by black boxes.

CRISPR repeat sequences differ between CRISPR-Cas sub-types and efficient adaptation is dependent on the repeat sequence^16,62^. Low resolution structural data suggest the type I-E repeat forms a single bend in the center of the CRISPR repeat binding channel^17,33,62^. Swapping the type I-F CRISPR repeat for the I-E CRISPR repeat does not impact the first transesterification reaction, but reduces the efficiency of the second transesterification, despite similar sequence lengths^43^. To understand if there were any conserved sequence features of the I-F repeat that might explain why the second transesterification reaction efficiency is reduced or outright abolished by swapping the repeat sequences, 2,804 type I-F repeats were aligned and plotted as a logo (**Figure 5E).** This analysis revealed that the 28 bp type I-F repeat is well conserved, including four purine-pyrimidine steps located in the inverted repeats (**Figure 5F**). The position of the DNA distortions observed in the repeat channel aligns well with these conserved purine-pyrimidine steps, which are particularly susceptible to DNA kinking (**Figure 5C-F**)^63^. Together with previous biochemical and mutagenesis studies^43^, these findings reveal that the repeat sequence is subject to sub-type specific distortions that function to further enhance the specificity of the CRISPR integrase. These DNA distortions ensure the second transesterification reaction occurs only when the integrase docks onto the correct repeat.

## DISCUSSION

Previous studies of CRISPR integrases have identified how genome folding is necessary for site-specific integration of foreign DNA^17,43^. However, the mechanisms of integrase activation and regulation remain poorly understood. Here we determine seven structures of the type I-F CRISPR integrase that collectively explain how auto-inhibition and allosteric activation regulate the CRISPR integrase. Cas1-2/3 forms a heterohexameric complex with a positively charged face that binds short fragments of foreign DNA (**Figure 1**). In this DNA capture state, the HD domain of Cas3 is blocked by a loop from the RecA1 domain (**Figure 2**). Binding to short fragments of DNA triggers conformational rearrangements that expose additional DNA interaction sites (**Figure 3**). Full integration of foreign DNA via sequential Cas1-mediated transesterification reactions on either side of the complex only occurs in the absence of a PAM (**Figure 4**). During full integration, the CRISPR repeat channel distorts the B-form repeat to direct the repeat DNA into the second transesterification site (**Figure 5**). Thus, these seven volumes and two atomic models provide step-wise snapshots of CRISPR adaptation.

These snapshots identify a key structural checkpoint during CRISPR adaptation (**Figure 1**, and **3**). Cas3 blocks Cas2 association with the folded CRISPR leader (**Figure 1**). After foreign DNA binding, Cas3 is displaced to expose two CRISPR leader binding sites on either side of Cas2 (**Figure 3**). DNA binding motifs upstream of CRISPR arrays^19,64^ correlate with the observed diversity of CRISPR adaptation strategies in bacteria and archaea^17,19,43,64–67^. This observation suggests that recognition of specific genome structures by the integrase is a conserved feature of CRISPR adaptation. Our study shows that foreign DNA allosterically regulates Cas1-2/3 integrase conformation and is sufficient to expose DNA binding sites necessary for homing the integrase to the CRISPR.

Previous work has shown that nuclease/helicase-competent Cas3 is involved in both naïve and primed acquisition of new foreign DNA sequences into the CRISPR array^21,68,69^. This study reveals a direct structural role for Cas3 in regulating new sequence acquisition in the type I-F system. Cas3 directly gates access to key DNA binding sites on the type I-F integrase (**Figure 1**, **Figure 2**). To our knowledge, interactions between Cas1 and Cas3 have only been tested in type I-F system. However, Cas1-2 complexes have been shown to attenuate Cas3 nuclease activity and facilitate long-range translocation, revealing that Cas1-2 integrases interact with Cas3 even in the absence of genetic fusion^38^. To determine if Cas3 might regulate integrases beyond the I-F system, we used AlphaFold3 to predict structures of Cas1-2 heterohexamer from the type I-E system of *E. coli* in the presence of two Cas3 molecules, both with and without foreign DNA fragments (**Figure S13)**. AlphaFold3 consistently predicted nearly identical Cas1- Cas3 interactions across five different predicted structures(**Figure S13**). In all cases, Cas3 obstructs the face of Cas1 that is required to interact with IHF and the CRISPR leader during integration in the type I-E system (**Figure S13**)^17^. These predictions suggest that Cas3 and Cas1 may function together to regulate adaptation in other type I CRISPR systems. Our study did not consider the effects of helicase or nuclease mutations on gating the DNA binding sites of the Cas1-2/3 integrase, nor did we attempt to biochemically characterize Cas1-Cas3 interactions outside of the type I-F system. However, together these observations guide future work aimed at understanding the functional and physical link between Cas3 and the integrase in type I CRISPR systems^38^

Cas3 is a superfamily 2 (SF2) helicase^34^. Helicases are critical components of genome replication and repair, but they also play important roles in resolving nucleic acid structures and rearranging chromatin to regulate gene expression^71,72^. Phylogenetically, Cas3 proteins are most closely related to three families of eukaryotic helicases: DEAH/RHA SF2 helicases that remodel RNA structure^73,74^, NS3 helicases found in eukaryotic RNA viruses^75,76^,and eukaryotic Snf2-type motor ATPases that power chromatin remodelling^34,72^. IHF introduces dramatic bends during DNA mobilization^17,43,77,78^, but also maintains bacterial chromosome architecture^79,80^, presumably through the same kinking mechanism. The association of Cas3 with the folded genome and the close phylogenetic relationship to chromatin and RNA structure remodelers raises the intriguing possibility that Cas3 may have a secondary function re-arranging bacterial chromatin in a process that is analogous to eukaryotic chromatin remodeling.

Another intriguing parallel raised by this study and other recent studies of CRISPR integration^17,43^ is the requirement of DNA binding and bending proteins to integrate foreign DNA. In eukaryotes and archaea, nucleosomes are essential structures that regulate genome architecture and gene expression^81–83^. The retroviral integrase from human immunodeficiency virus (HIV) interacts with histones and integrates viral DNA adjacent to nucleosomes^84,85^. R2 retrotransposons forego an integrase and instead use a combination of an endonuclease and a target-primed reverse transcriptase to insert foreign DNA into nucleosome-bound DNA^86^. These findings suggest that genetic parasites across the domains of life target host DNA bending proteins to facilitate integration.

Targeted delivery of foreign nucleic acids is an important step for developing gene editing and molecular recording tools that precisely paste foreign sequences at new locations in the genome^87–90^. Recent studies using CRISPR-associated transposons or endonuclease/target-primed reverse transcriptases for genome editing reveal that integration efficiency remains a major barrier to genome editing^88,91^. Though *in vitro* evolution experiments have dramatically improved this efficiency^92^, current genome editing tools are not designed to mimic integrases that target three dimensional structures within target genomes. To our knowledge, current genome editing tools lack sophisticated mechanisms of allosteric activation. This study observes the allosteric activation of the CRISPR integrase by the foreign DNA payload to recognize a three-dimensional structure in a bacterial genome, providing two important mechanistic insights into integrase function. This functionality should be leveraged in future studies to explore new methods of safe and efficient target insertion at safe harbor sites already encoded in the genome.

## MATERIALS & METHODS

### Protein purification

*P. aeruginosa* Cas1-2/3 heterohexamer complexes were purified as previously described^19^. Briefly, StrepII-tagged Cas1 (WP_003139228.1) and untagged Cas2/3 (WP_003139227.1) encoded on a single plasmid were overexpressed in *E. coli* BL21(DE3) (Addgene #89240). Cell pellets were lysed via sonication in Cas1-2/3 lysis buffer (50 mM HEPES pH 7.5, 500 mM KCl, 10% Glycerol, 1 mM DTT), supplemented with 0.3x Halt Protease Inhibitor Cocktail (Thermo Fisher Scientific), at 4 °C. Lysate was clarifed as above. StrepII-tagged Cas1-Cas2/3 complexes were affinity purified on StrepTrap HP resin (GE Healthcare) and eluted with Cas1-2/3 lysis buffer containing 3 mM desthiobiotin (Sigma-Aldrich). Eluate was concentrated at 4 °C (Corning Spin-X concentrators), before purification over a Superdex 200 size-exclusion column (Cytiva) equilibrated in 10 mM HEPES pH 7.5, 500 mM Potassium Glutamate, and 10% Glycerol.

*P. aeruginosa* IHF heterodimer was purified as previously described^19^. Briefly, 6xHis-tagged IHFα and StrepII-tagged IHFβ were co-expressed in *E. coli* BL21(DE3) (Addgene #149384, #149385). Cell pellets were lysed by sonication in IHF lysis buffer (25 mM HEPES-NaOH pH 7.5, 500 mM NaCl, 10 mM Imidazole, 1 mM TCEP, 5% Glycerol), supplemented with 0.3x Halt Protease Inhibitor Cocktail (Thermo Fisher Scientific), at 4 °C. Lysate was clarified by two rounds of centrifugation at 10,000 x g for 15 minutes, at 4 °C. His-tagged IHF was captured on HisTrap HP resin (Cytiva), and eluted with 500 mM Imidazole. Affinity tags were cleaved using PreScision protease, and the PreScision protease and remaining 6x-His-IHFα were removed by affinity chromatography using HisTrap HP resin (Cytiva). Untagged IHF heterodimer was then further purified on Heparin Sepharose (Cytiva) and eluted with a linear gradient to a buffer containing 2 M NaCl. Fractions containing IHF heterodimer were concentrated and further purified by Size Exclusion Chromatography (SEC) on a Superdex 75 column (Cytiva) equilibrated in IHF buffer (25 mM HEPES-NaOH pH 7.5, 200 mM NaCl, 5 % Glycerol).

### Cryo-EM sample preparation and data acquisition of the Cas1-2/3 capture complex

Frozen aliquots of purified Cas1-2/3 complex (12.8 μM) were shipped on dry ice to the National Center for CryoEM Access and Training (NCCAT) and the Simons Electron Microscopy Center located at the New York Structural Biology Center (NYSBC). 3 µL of sample was spotted on glow discharged Quantifoil R0.6/1 Cu 300 mesh grids (Quantifoil Mico Tools GmbH) and blotted for 2 seconds at 10 °C and 100% humidity using a Leica EM-GP2 (Leica Microsystems) prior to plunge freezing in liquid ethane. The dataset was collected on a Titan Krios (Thermo Fisher Scientific) at NYSBC operating at 300 keV with a Gatan K3 direct electron detector and a GIF-Quantum energy filter with a 20 eV slit width at a nominal magnification of 81,000x. 40 frame movies were collected at a 50 milliseconds frame rate and a dose rate of 25.26 e^-^/Å^2^/s for a total dose of 52.51 e^-^/Å^2^ with a pixel size of 1.0691 Å/px over a defocus range of -0.8 to -2.5 μm.

Holes were identified and targeted and automated data collection was carried out using Leginon^93^. Preliminary dataprocessing was done on the fly in cryoSPARC Live^61^.

### Conservation analysis of Cas1 and Cas2/3

We used two complementary approaches to measure conservation of residues in Cas1 and Cas2/3 proteins. To map conserved residues in the structure, we use CasFinder to identify unique occurrences of type I-F CRISPR systems in NCBI’s database of complete bacterial and archaeal genomes^94^. Complete genomes from NCBI were downloaded on July 11, 2023 and open-reading frames annotated using Prodigal.^95^ These open reading frames were then queried for *cas* genes using MacsyFinder v1.0.5^96^ with the following parameters: *“macsyfinder -- sequence-db <OPEN_READING_FRAMES> --db-type gembase -d <CRISPR_SUBTYPE_DEFINITIONS> -p <HMM_PROFILES> -w 50 -vv all”*. HMM profiles and classification definitions (*CRISPR_subtype_definitions*) used in MacsyFinder are acquired from the edited version of CasFinder v3.1.0^94^ to include definitions for the CAST systems (available at https://github.com/macsy-models/CasFinder). Type I-F Cas2/3 sequences (any sequence annotated with "cas3f","cas3","cas3HD", n=1922) and Cas1 sequences (n=1838) were extracted from the database of Prodigal-annotated open-reading frames and identical sequences removed with CD-hit (-c 1)^95,97^. The remaining 739 (Cas1) or 970 (Cas2/3) sequences were aligned in MAFFT (mafft --genafpair --maxiterate 1000), and poorly aligned sequences were removed using MaxAlign (flags: -v=1, -a). The resulting alignments of 679 (Cas1) and 745 (Cas2/3) sequences were submitted to Consurf^98^ with the Cas1-2/3 atomic model to calculate conservation scores. PA14 Cas1 and Cas2/3 sequences (from genome NC_008463) were associated with the submitted atomic model. Conservation scores were mapped onto atomic models using ChimeraX (**Figure S4, S9, S12**)^99^.

To understand if the global architecture of the Cas1-2/3 complex was conserved, the Cas2/3 sequence from *Pseudomonas aeruginosa* strain UCBPP-PA14 (100% identical to multi-species accession WP_003139227.1) was used to query the nr_70_Mar12 database using the ProtBLAST/PSI-BLAST tool on the MPI Bioinformatics Toolkit server (Scoring Matrix: BLOSUM62; E-value cutoff for reporting: 1e-3; E-Value cutoff for inclusion: 1e-6; Max target hits: 1000).^100^ The resulting alignment of 1000 sequences was downloaded and 170 sequences missing Cas2 or Cas3 domains were manually removed. MaxAlign was used to remove 145 additional poorly aligned sequences (flags: (-v=1, -a).^101^ Gappy columns were removed using trimAL (flags: -gt 0.7) to generate an alignment of Cas2/3 across 1071 positions.^102^ This alignment was used to build a phylogenetic tree using Fasttree (flags: -pseudo, -wag, -gamma), which was visualized in R using ggtree (**Figure S5**)^103,104^.

### Assembly and purification of the delivery complex

Foreign DNA fragments (32-34 bp) were ordered from IDT as 100 nmole ssDNA oligonucleotides and resuspended to a working concentration of 100 uM in 1x TE buffer. To generate dsDNA foreign DNA fragments, 2 nmole of ssDNA was suspended in 0.5X hybridization buffer (20 mM Tris-HCl pH 7.5, 220 mM K-Glutamate, 5 mM EDTA, 1 mM TCEP, 2 % glycerol) and incubated at 95 °C for 5 minutes, followed by 25 °C for 10 minutes. Samples were then denatured at 100 °C for 5 minutes, and slow cooled to 25 °C at a rate of 6 degrees/5 minutes for an hour, to a final temperature of 25 °C.

A 1:1 mixture of purified Cas1-2/3 and dsDNA was created by mixing 2 nmoles of purified Cas1-2/3 thawed on ice then warmed to 25 °C with 2 nmoles of annealed dsDNA containing a single 3’ GG extension. This mixture was incubated at 25 °C for 20 minutes, then centrifuged at 4 °C, 22000 g for 20 minutes to pellet any aggregate. The supernatant was collected in a 1 mL syringe and injected onto a Superdex 200 10/300 column (Cytiva) equilibrated with 20 mM Tris-HCl pH 7.5, 200 mM monopotassium glutamate, 5 mM EDTA, 1 mM TCEP, 2 % Glycerol.

Fractions (0.25 ml) were individually concentrated using 100 kDa molecular weight cuttoff concentrator at 4 °C, 15000 g, to ∼20 uL. All fractions were analyzed on SDS-PAGE gels (15 % resolving) to determine protein composition and 8 % Urea-PAGE gels to determine nucleic acid composition. The fifth SEC fraction contained all DNAs and proteins of interest and was further analyzed by cryo-EM.

### Cryo-EM sample preparation and data acquisition of the Cas1-2/3 delivery complex

Frozen aliquots of purified Cas1-2/3 delivery complex bound to foreign DNA (45.4 μM) were shipped on dry ice to the National Center for CryoEM Access and Training (NCCAT) and the Simons Electron Microscopy Center located at the New York Structural Biology Center (NYSBC). Sample was thawed and diluted 1:10 in SEC buffer to a concentration of 4.54 μM. 3 μ L of diluted sample was spotted on glow discharged Quantifoil R1.2/1.3 Cu 300 mesh grids (Quantifoil Micro Tools GmbH), blotted for 4.5 seconds at force 0 and 4 °C, 100% humidity, and plunge frozen in liquid ethane using a Vitrobot (Mk. IV, Thermo Fisher Scientific). The dataset was collected on a Titan Krios (Thermo Fisher Scientific) at NYSBC operating at 300 keV with a Falcon 4EC direct electron detector at a nominal magnification of 96,000x. 40 frame movies were collected at a 150 ms frame rate and a dose rate of 8.63 e^-^/Å^2^/s for a total dose of 51.78 e^-^/Å^2^ with a pixel size of 0.833 Å/px over a defocus range of -0.9 to -2.8 um. Holes were identified and targeted and automated data collection was carried out using Leginon^92^.Preliminary dataprocessing was done on the fly in cryoSPARC Live^61^.

### Assembly and purification of the integration complex

To assemble a 200 bp fragment of the CRISPR leader and first repeat and spacer, 4 nmol of 200 bp ssDNA fragments were ordered from IDT and suspended to 200 uM in a modified hybridization buffer (20 mM HEPES pH 7.5, 200 mM K-glutamate, 2% glycerol, 1 mM TCEP). ssDNA fragments were combined in a 1:1 ratio to generate 100 uM dsDNA, heated to 100 °C for 5 minutes, and then slow-cooled to 25 °C over the course of an hour. Annealed oligos were stored at -80 °C. 3.75 nmoles of purified IHF (at 1234 uM) was diluted to a final volume of 125 uL (30 uM) in SEC buffer (20 mM HEPS pH 7.5, 200 mM K-Glutamate, 5 mM MnCl_2_,7.5 mM spermidine, 2% glycerol, 1 mM TCEP) at 25 °C to avoid precipitation of IHF. 1.5 nmoles of annealed CRISPR fragment was diluted to a final volume of 95 uL (15 uM) in the same SEC buffer. Both the pre-diluted IHF solution and pre-diluted CRISPR solution were briefly warmed to 25 °C prior to mixing (220 uL final volume) and incubation for 25 °C for 10 minutes.

1.5 nmoles of purified Cas1-2/3 was mixed with 1.875 nmoles of foreign DNA fragment and 1.4 uL of 1 M MnCl_2_ and 2.1 uL of 1 M spermidine in pH neutralized buffer and incubated for 10 minutes at 25 °C.

The CRISPR-IHF reaction was mixed with the Cas1-2/3-foreign DNA reaction and incubated for 20 minutes at 35 °C. Precipitates were removed by centrifugation at 4 °C, 20,000xg, prior to size exclusion. A Superose 6 10/300 column was equilibrated to SEC buffer (20 mM HEPS pH 7.5, 200 mM K-Glutamate, 5 mM MnCl_2_, 7.5 mM spermidine, 2% glycerol, 1 mM TCEP) at 4 °C. 0.5 mL fractions were collected, analyzed on SDS-PAGE (15 %) and UREA-PAGE (8 %) gels and concentrated to 10 uL (∼4 uM for PAM, ∼3 uM for trimmed).

### Cryo-EM sample preparation and data acquisition of the Cas1-2/3 integration complex

Purified integration complex with either 34 bp of dsDNA that included a ‘GG’ PAM or 32 bp dsDNA fragment with no PAM was diluted to a concentration of ∼1 µM in SEC buffer lacking glycerol (20 mM HEPS pH 7.5, 200 mM K-Glutamate, 5 mM MnCl_2_ 7.5 mM spermidine, 2 % glycerol, 1 mM TCEP). 3 µl of diluted integration complex was spotted on Quantifoil Au 300 R2/1 + 2 nm continuous C grids (Quantifoil Micro Tools GmbH) that were glow discharged for 45 seconds with a 10 second hold (easiGlow, Pelco). The grids were then blotted for 5 seconds without incubation, with a blot force of "5", at 100% humidity and 4 °C followed by plunge freezing using a Vitrobot (Mk. IV, Thermo Fisher Scientific). The datasets for both the trimmed and PAM-containing integration complexes were collected on Montana State University’s Talos Arctica transmission electron microscope (Thermo Fisher Scientific), with a field emission gun operating at an accelerating voltage of 200 keV using parallel illumination conditions. Movies were acquired using a Gatan K3 direct electron detector, operated in electron counting mode targeting a total electron exposure of 65.67 e-/Å^2^ over 50 frames (5.992 second exposure, 0.12 second frame time). The SerialEM data collection software was used to collect micrographs at 36,000x nominal magnification (1.152 Å/pixel at the specimen level) over a defocus range of -0.5 µm to -2.0 µm. Stage movement was used to target the center of four 2.0 µm holes for focusing, and image shift was used to acquire high magnification images in the center of each of the holes. Preliminary data processing was done on the fly in cryoSPARC Live.

### Cryo-EM image processing

Movies were motion and CTF-corrected in cryoSPARC using patch motion and patch CTF correction^61^. For details, see Supplementary Figure 1 (capture complex), Supplementary Figure 7 (delivery complex), Supplementary Figure 10 (integration complex with PAM-containing foreign DNA), and Supplementary Figure 11 (integration complex with foreign DNA, no PAM). In brief, initial rounds of blob picking (capture/delivery) or template picking (integration complexes) were used to generate 3D volumes from the dataset for a second round of template picking.

Template-picked particles were subject to multiple rounds of multi-class *ab initio* reconstructions and heterogenous refinements to identify the major 3D classes of the complex. Particle stacks corresponding to individual complex were further refined through iterative multi-class *ab initio* reconstructions and heterogeneous refinements, local refinement, 3D classification, and 3D variability analysis in cryoSPARC.

### Model building and validation

AlphaFold3 was used to generate a prelimnary model of the capture complex^70^. However, AlphaFold3’s predicted positioning of Cas3 does not correlate well with cryo-EM density maps for the capture and integration complexes, so in these instances, individual protein molecules from Alphafold were sequentially docked into the density. Protein and DNA segments were individually rigid-body fit into the EM density map using the “fit <MODEL> inMap <MAP>” command in UCSF ChimeraX^99^. Isolde was intitially used to morph the rigid body-fit initial model into the density map, followed by Real Space Refinement with morphing (no secondary structure restraints, ignore symmetry conflicts) in Phenix^44,46^. The resulting map and model were re-opened in Isolde/ChimeraX and any gross conformational issues were corrected with the Isolde simulation running. After a second round of Real Space refinement (no morphing, no secondary structure restraints), problem areas were inspected in Coot and restrained to ideal geometry, using secondary structure and German McClure distance restraints generated in ProSMART^105^. MolProbity and the PDB validation service server (https://validate-rcsb-1.wwpdb.org/) were used to identify problem regions subsequently corrected in Coot^106,107^.

Contacts and hydrogen bonds between residues were identified by ChimeraX v1.9 using the “contacts” and “hbonds” commands respectively, with default parameters.

### Conservation analysis of type I-F repeats

Complete bacterial (n=39,277) and archaeal (n=556) genomes and chromosomes were downloaded from the NCBI RefSeq Assembly database (accessed on July 11, 2023). CRISPR loci within 93,671 genomic and plasmid sequences were identified using the default parameters in CRISPRDetect v3^108^, which resulted in 37,477 high-confidence CRISPR loci predictions (array quality score > 3). Type I-F repeat sequences were extracted from the default output file of CRISPR repeat sequences and aligned in MAFFT. Logo plots were generated using Weblogo^109^.

## ACKNOWLEDGEMENTS

We thank members of the Wiedenheft lab, and particularly Nathaniel Burman, Royce Wilkinson, and Murat Buyukyoruk, for their invaluable input and discussions. Research in the Wiedenheft laboratory is supported by the National Institutes of Health (R35GM134867), the M. J. Murdock Charitable Trust, and the Montana Agricultural Experimental Station. A.S-F. is supported by the National Institutes of Health (K99GM147842, R00GM147842), and by the Postdoctoral Enrichment Program Award from the Burroughs Wellcome Fund (G-1021106.01). Some of the microscopy was performed using resources provided by the National Center for CryoEM Access and Training (NCCAT) and the Simons Electron Microscopy Center located at the New York Structural Biology Center, supported by the NIH Common Fund Transformative High Resolution Cryo-Electron Microscopy program (U24 GM129539, and NIGMS R24 GM154192) and by grants from the Simons Foundation (SF349247) and NY State Assembly. Funding for the Montana State University cryo-EM Core Facility (RRID:SCR_026324) was contributed by National Science Foundation (DBI-1828765), the MJ Murdock Charitable Trust, The National Institute of General Medical Sciences (P30GM140963) and the MSU Office of Research, Economic Development and Graduate Education. Computational efforts were performed on the Tempest High Performance Computing System, operated and supported by University Information Technology Research Cyberinfrastructure (RRID:SCR_026229) at Montana State University.

## DATA AVAILABILITY

Electron density maps are available at EMD-71091 (Cas1-2/3), EMD-71097 (Cas1-2/3 and foreign DNA), EMD-71112 (Integration complex with PAM – CRISPR-leader associated), EMD- 71101 (Integration complex with PAM – first transesterification), EMD-71114 (Integration complex without the PAM – CRISPR leader associated), EMD-71115 (Integration complex without the PAM – first transesterification), EMD-71116 (Integration complex without the PAM – full repeat). Atomic models are available at the Protein Data Bank under accession codes: 9P11 (Cas1-2/3), 9P1D (Cas1-2/3 and foreign DNA). Alignments and Newick files used for conservation and phylogenetic analyses are available as supplementary data.

## Abbreviations

CRISPR: Clustered regularly interspaced short palindromic repeats
Cas: CRISPR-associated

## SUPPLEMENTAL FIGURES

**Figure S1.**
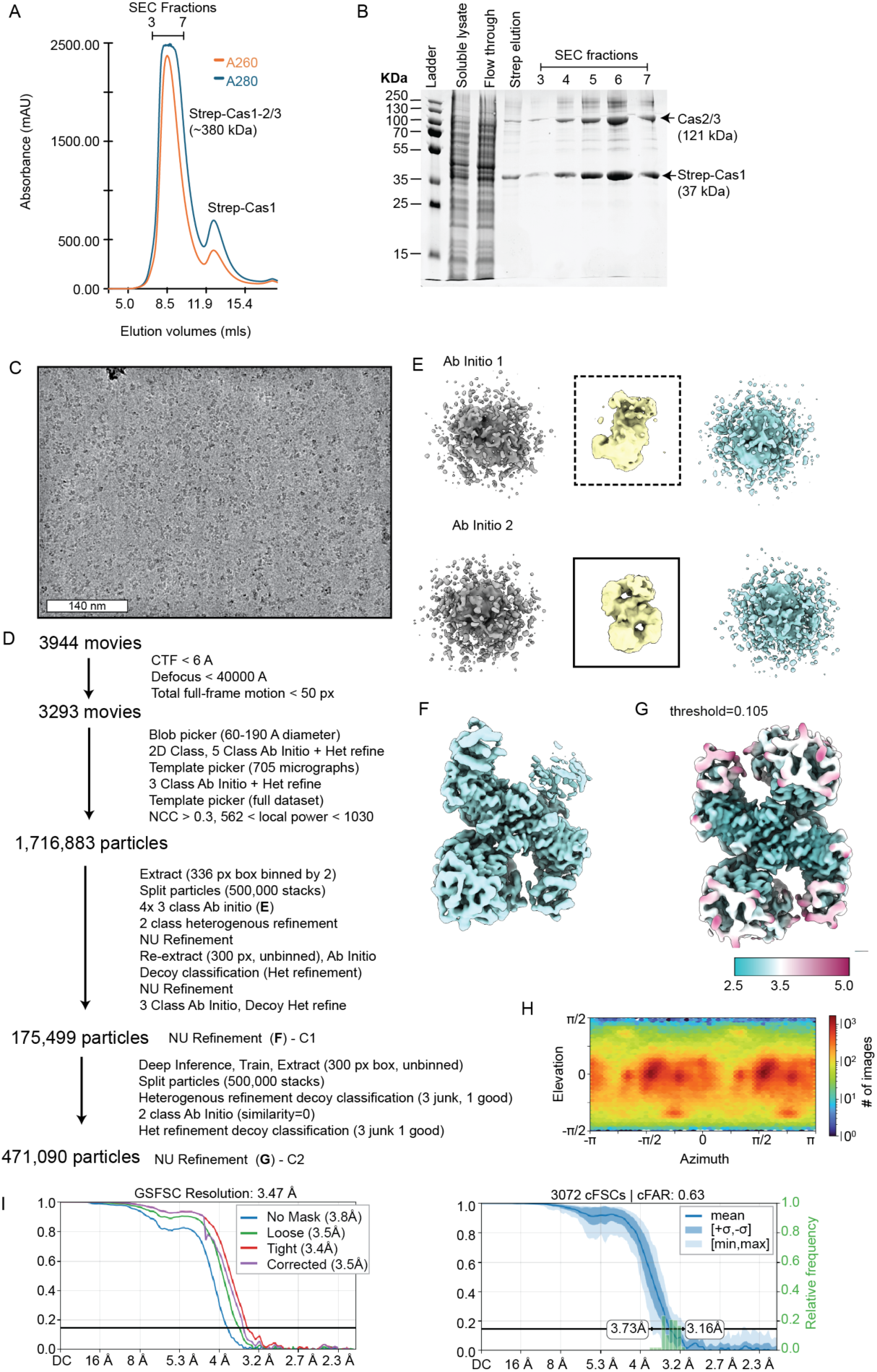
Purification and data processing of the Cas1-2/3 capture complex. Related to Figure 1. (A) Size exclusion chromatography (SEC) of Cas1-2/3 results in a monodispersed complex with an estimated molecular weight of 380 kDa. (B) SDS-PAGE of the fractions following affinity purification, and size exclusion. (C) Sample micrograph at a nominal magnification of 81,000. (D) Data processing summary. (E) Results from two of four total, 3-class *Ab initio* reconstructions and heterogenous refinements provided two volumes (boxed) for baited heterogeneous refinement (dashed box indicates volume with severe anisotropy, solid box indicates a volume with well-distributed views). (F) Non-uniform (NU) refinement of 175,499 particles used to train cryoSPARC’s Deep Inference job. (G) Final NU refinement of 471,090 particles reconstructed with C2 symmetry imposed (threshold = 0.105). Coloring by local resolution (estimation at FSC 0.5) reveals most of the complex is resolved below 3.5 Å. (H) Azimuth plot from cryoSPARC of the final C2 NU refinement. (I) FSC and conical FSC plots of the final C2 NU refinement.

**Figure S2.**
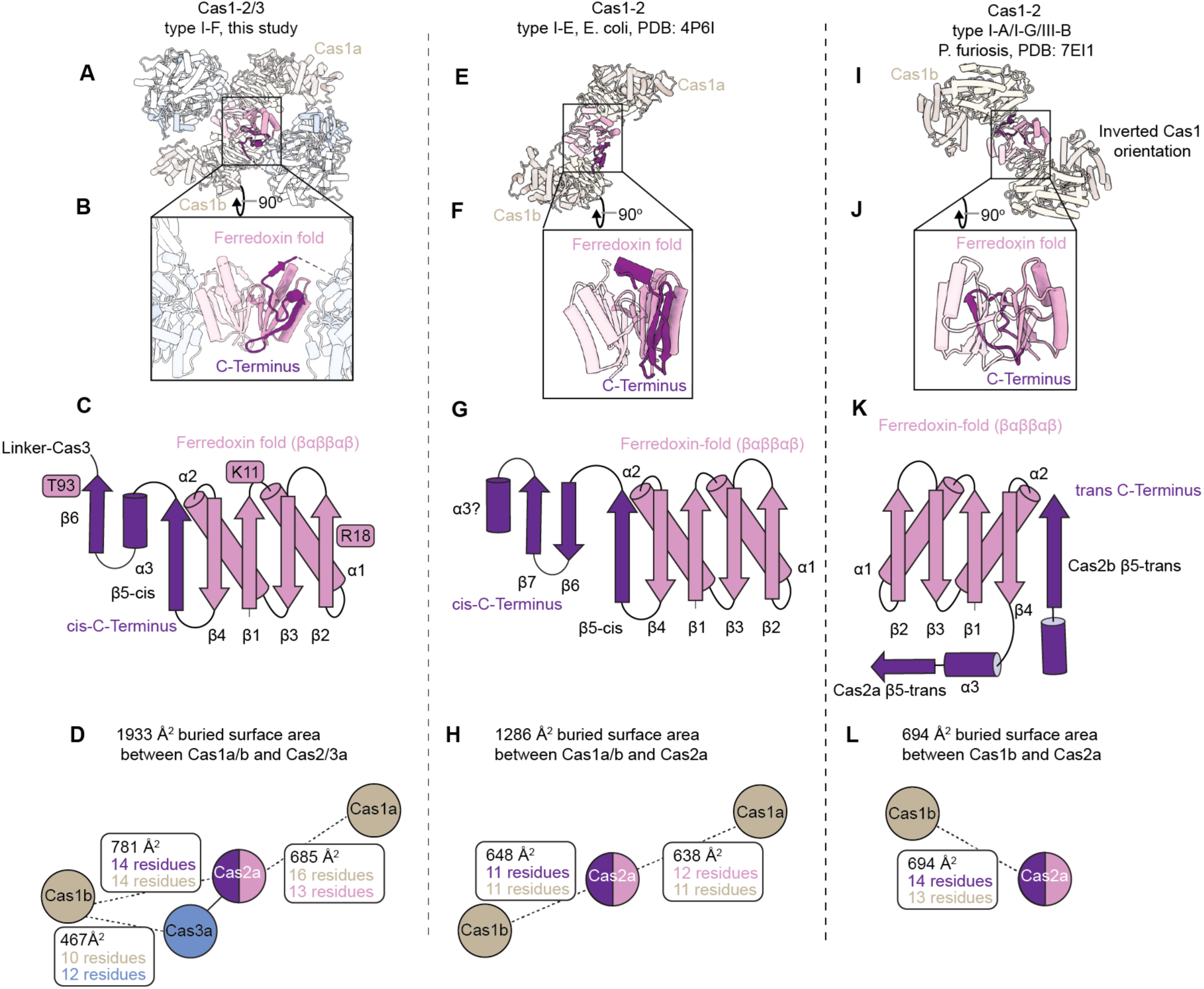
The C-terminal topology of Cas2 determines the Cas1 interface. Related to Figure 1. (A, E, I). The variable Cas2 C-terminus determines the identity and extent of partner interaction. Structures of experimentally determined Cas1-2 complexes from the PDB shown at 50% transparency, with one subunit of the Cas2 dimer colored to show the core ferredoxin-like fold (pink) and the variable C-terminus (purple). (B, F, J) Cas2-containing subunit only, rotated 90 degrees to visualize the dimer interface. (C, G, K) Ribbon diagrams for each Cas2 structure in panels B, F, J. (D, H, L) Interaction diagrams depicting the subunit interactions for a single Cas2 dimer, with the number of residues and buried surface area of each interface depicted.

**Figure S3.**
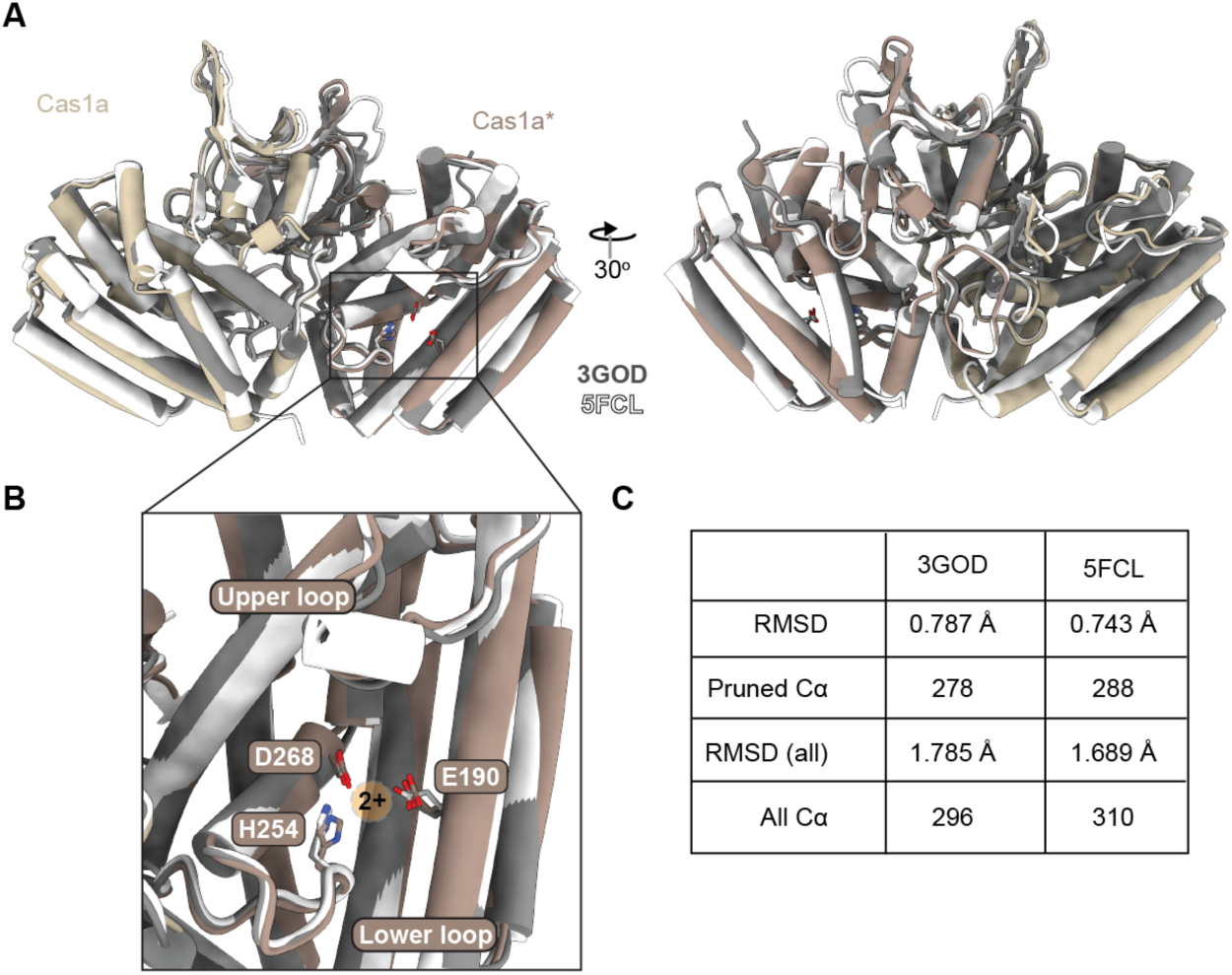
Cas2/3 association does not change Cas1 dimer architecture. Related to Figure 1. (A) Structural overlay of previously determined Cas1 dimer crystal structures (PDB: 3GOD, *P. aeruginosa* in grey, PDB: 5FCL, *P. atrosepticum* in white) onto one Cas1 dimer from the Cas1-2/3 complex (colored as in Fig 1). (B) Close-up of the Cas1* active site shows nearly perfect overlap of active site residues involved in coordinating a divalent metal ion (not observed in the cryo-EM structure due to EDTA in the purification buffer) (C) Table of root mean square deviations across pruned and unpruned Cα atoms, from the matchmaker (mmaker) command in UCSF ChimeraX.

**Figure S4.**
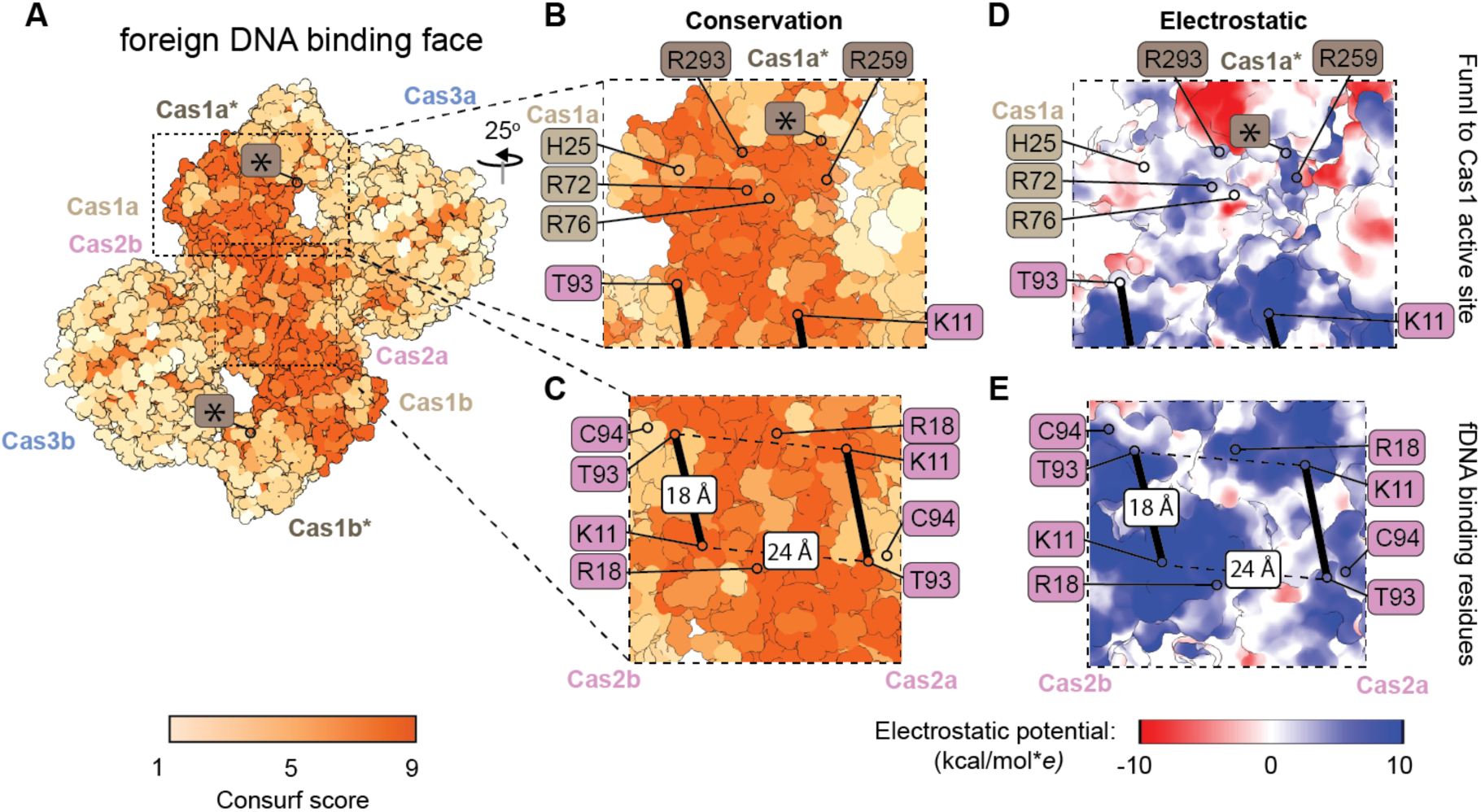
Positively charged foreign DNA binding residues are conserved in the Cas1-2/3 complex. Related to Figure 1. (A) Multiple sequence alignments of Cas1 (n=679) and Cas2/3 (n=745) from type I-F systems were used to calculate Consurf conservation scores, and mapped back to the atomic model of the complex (1 – least conserved, 9 – most conserved). Asterisk (*) indicates the Cas1 transesterification sites. (B, D). Conserved positively charged residues line a channel formed from the foreign DNA binding face to the Cas1 active site. Channel edges indicated with thick black lines. Distances are measured between the T93 OG1 and K11NZ using the distances command in ChimeraX. (C, E) DNA binding residues on the foreign DNA binding face of Cas2 sit in conserved patches of positive charge. Channel edges indicated with thick black lines.

**Figure S5.**
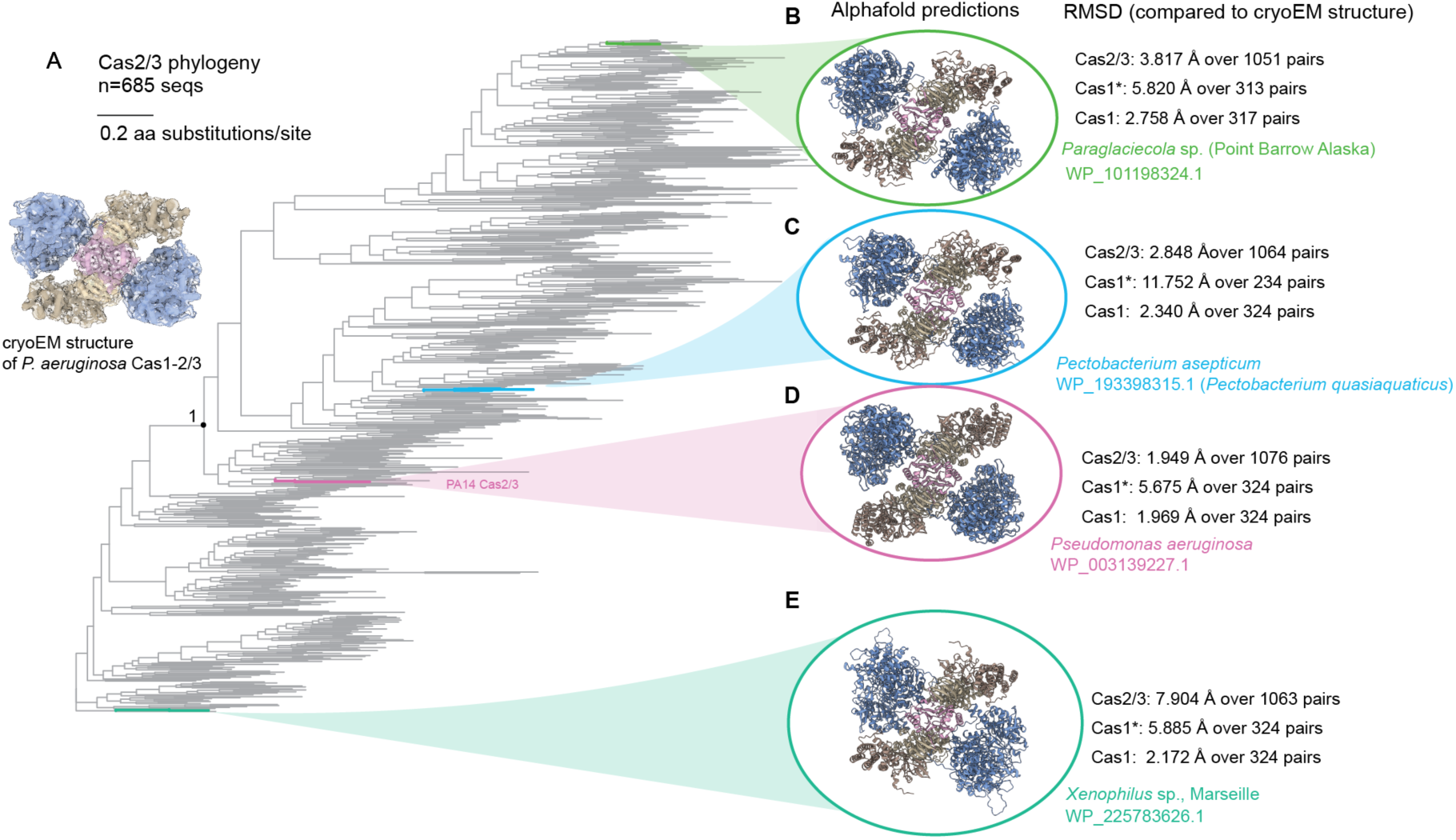
The four-bladed propeller shape of Cas1-2/3 systems is conserved. Related to Figure 1. (A) Phylogenetic tree of 685 Cas2/3 sequences. Sequences were identified through three iterations of PSI-BLAST, aligned in MAFFT, and the phylogenetic tree built in FastTree (see Methods). (B-E). AlphaFold3 predictions of Cas1-Cas2/3 heterohexamers reveal structural conservation. AlphaFold3 prediction were superimposed on the experimentally-determined structure using the mmaker command in ChimeraX. Root-mean square deviation of each subunit compared to across diverse type I-F systems are reported.

**Figure S6.**
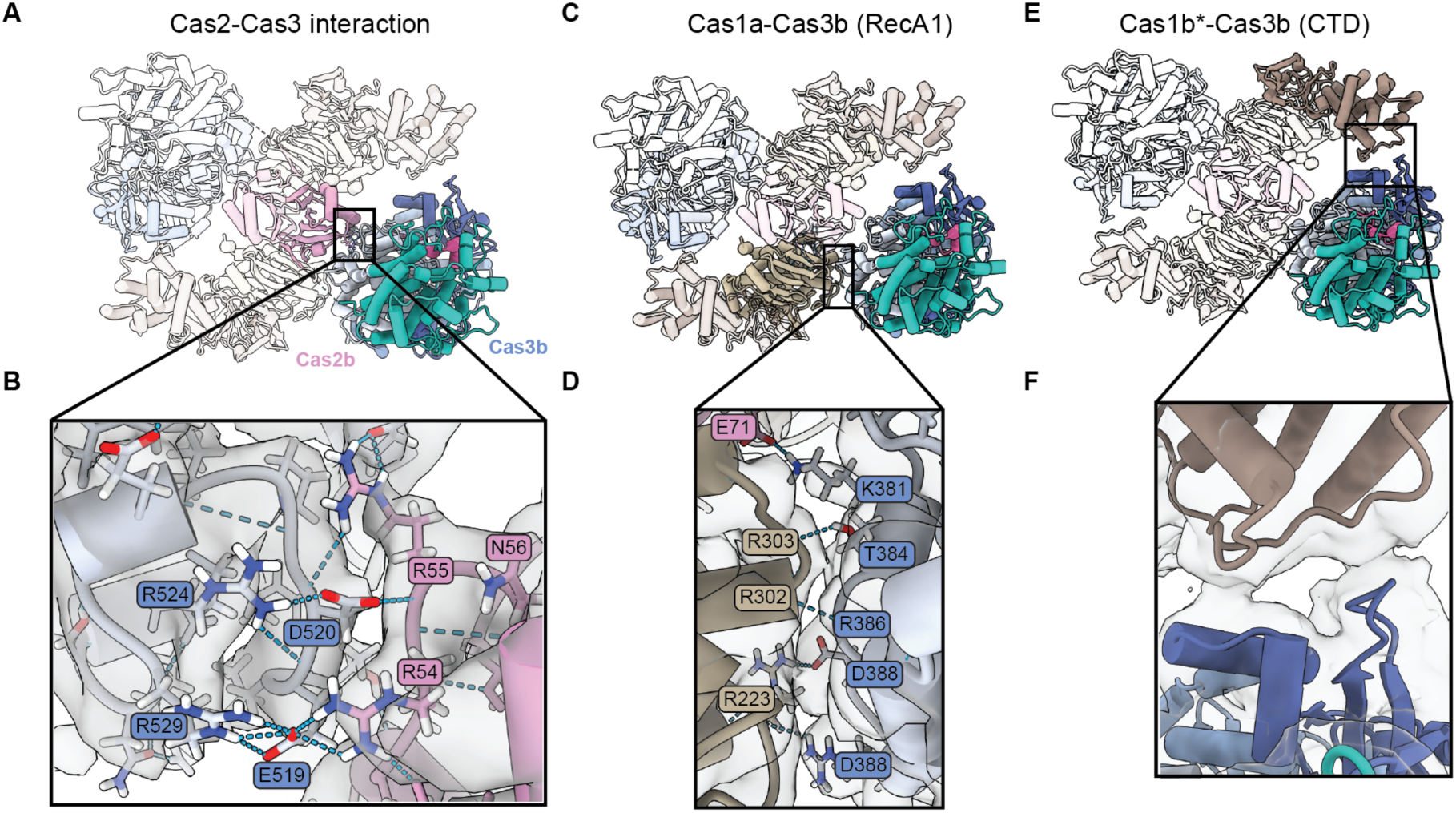
Three interfaces clamp Cas3 in place in the absence of DNA or Cascade. Related to Figure 1 and Figure 3. (A) Global view of the Cas1-2/3 complex, with the Cas2/3 interface responsible for blocking genome-interacting residues (R55) and locking the RecA1 domain in place shown for a single Cas2/3 subunit, all other subunits displayed at 90% transparency. (B) Hydrogen bond interactions between Cas2 (K50, R54, R55) and Cas3 (D472, D473, N373, E519 D520) reveal that the CRISPR leader interaction residues are sequestered in Cas3’s RecA1 domain, density map shown at 80%, contour level = 0.1. (C) Global view of the Cas1-2/3 complex, with the Cas1-Cas3 interface in the RecA1 domain highlighted in full color, all other subunits displayed at 90% transparency. (D) Side chain and main chain hydrogen bonds at the RecA1-Cas1 interface bury ∼500 Å^2^. Density map shown at 80%, contour level = 0.1. (E) Global view of the Cas1-2/3 complex, with the Cas1-Cas3 CTD domain highlighted in full color, all other subunits displayed at 90% transparency. (F) Flexibility at the Cas1-Cas3 CTD interface precludes side-chain modelling, likely due to multiple transient interaction between Cas1 and the CTD Density map shown at 80%, contour level = 0.0477. AlphaFold3 predicts multiple hydrogen bonds between acidic residues of Cas3 CTD and basic residues of Cas1.

**Figure S7.**
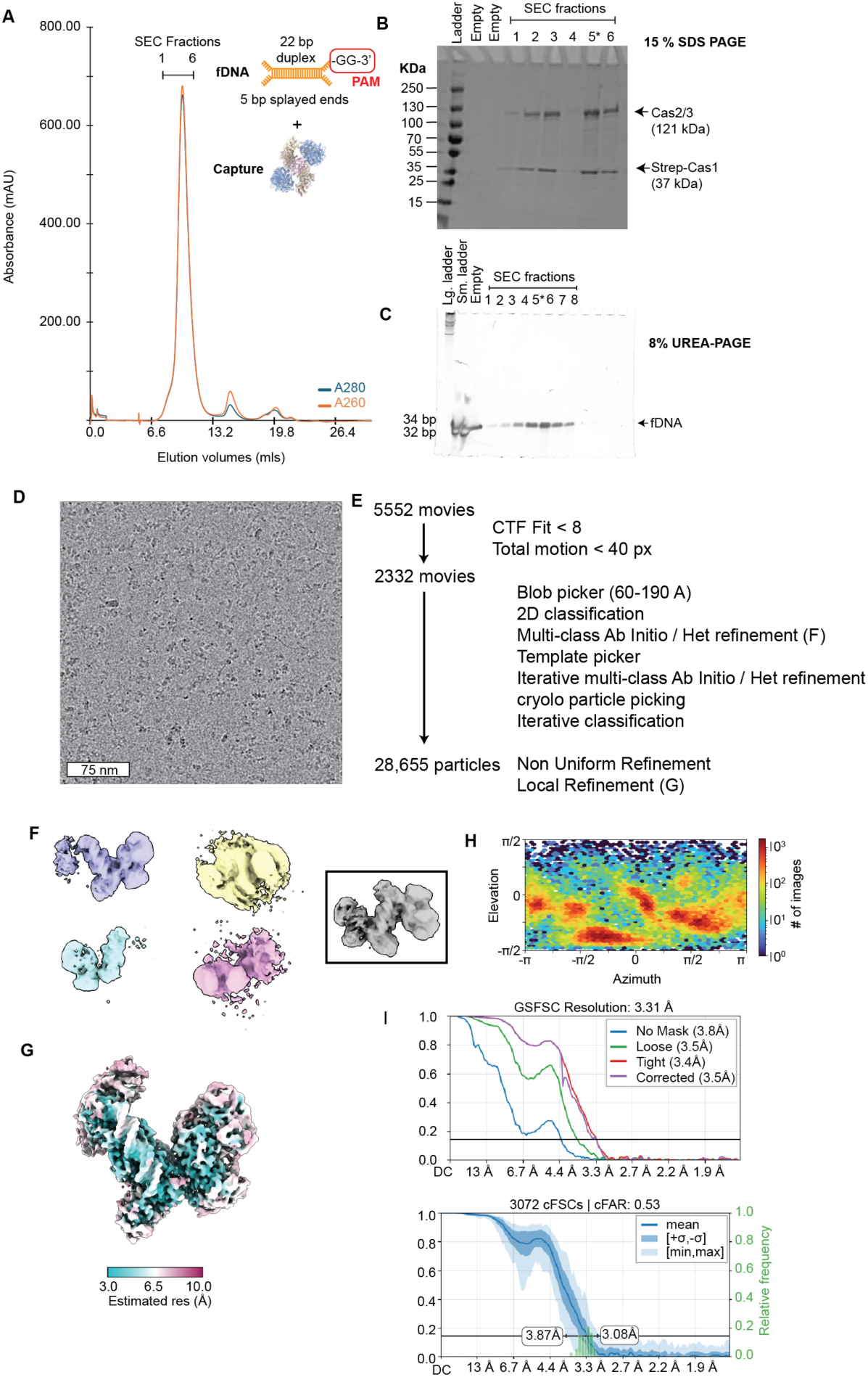
Purification and data processing of the Cas1-2/3 capture complex. Related to Figure 3. (A) The Cas1-2/3 complex bound to a fragment of DNA elutes from size exclusion chromatography as a monodispersed peak with an estimated molecular weight of 415 kDa. (B) SDS-PAGE stained with Coomassie blue reveals two proteins consistent with the sizes of Cas1 and Cas2/3. Asterisk (*) indicates fraction on which cryo-EM data was collected. Fraction four was lost due to spilled tube during preparation of SDS-PAGE gel (C) Urea-PAGE denaturing gel of SEC fractions confirms that the main peak contains a fragment of DNA. Asterisk (*) indicates fraction used for cryo-EM. (D) Sample micrograph at a nominal magnification of 96,000x. (E) Data processing summary. (F) Volumes from a 5-class *ab initio* reconstruction and heterogenous refinement after preliminary particle picking. The grey volume (boxed in black outline) was used subsequently as a template for template picking. (G) Local refinement of 28,655 particles colored by local resolution at estimation (FSC 0.5) reveals most of the complex is resolved below 6.5 Å but the edges of the complex are low resolution. (H) Azimuth plot reveals patchwork orientation bias, (I) FSC and conical FSC curves from cryoSPARC.

**Figure S8.**
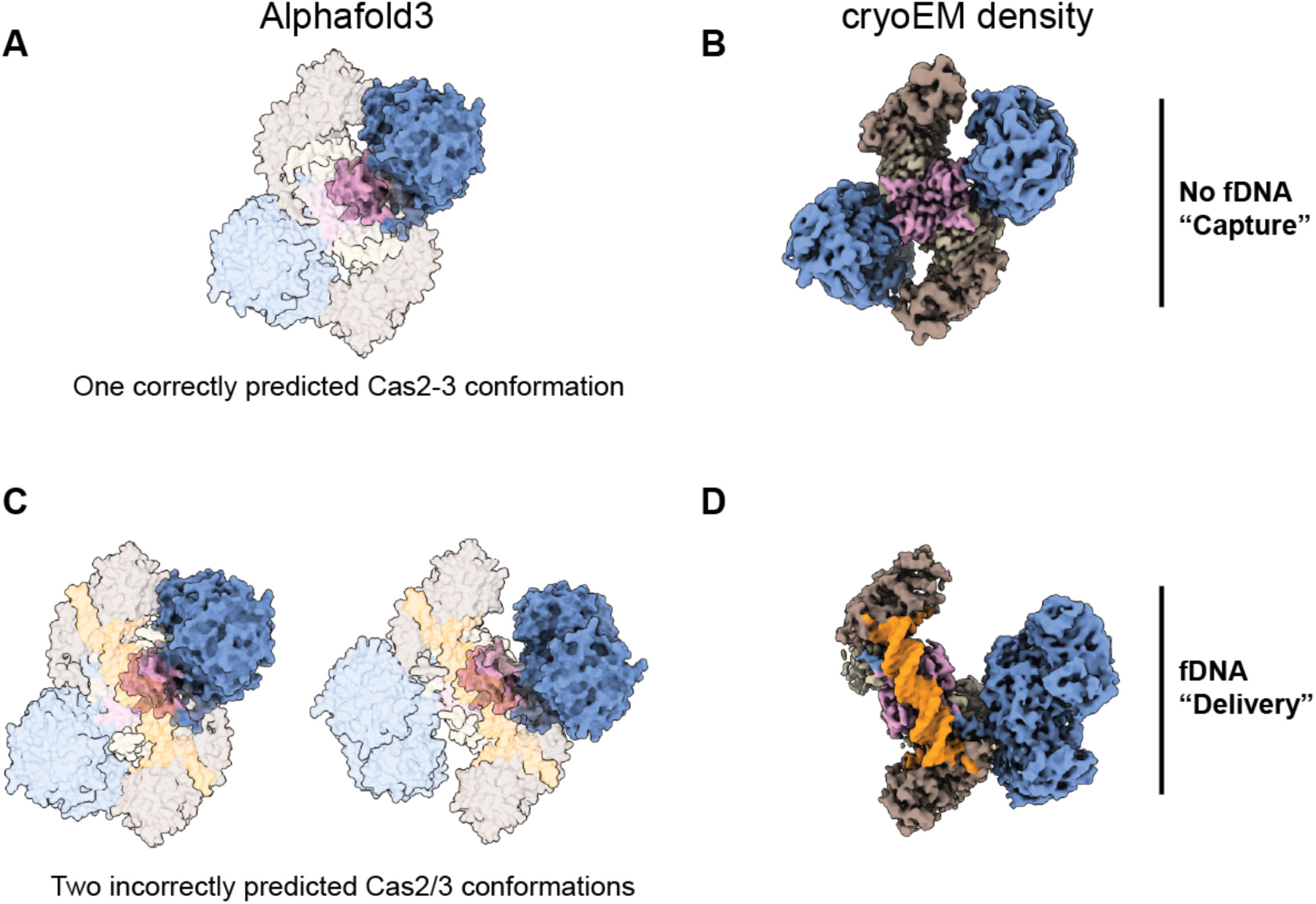
AlphaFold3 prediction support variable positioning of Cas3 after foreign DNA binding. Related to Figure 3. (A) AlphaFold3 prediction of the Cas1-2/3 complex bound in the absence of foreign DNA. (B) cryo-EM density for the Cas1-2/3 complex in the absence of foreign DNA agrees with AlphaFold3 prediction (map threshold=0.123). (C) AlphaFold3 predicts Cas3 in two distinct positions in the presence of a 32 bp foreign DNA fragment. (D) cryo-EM map of Cas1-2/3 bound to foreign DNA reveals one Cas3 is too flexible to resolve, while the other Cas3 domain is in an orientation that does not agree with the AlphaFold3 position (map threshold=0.195).

**Figure S9.**
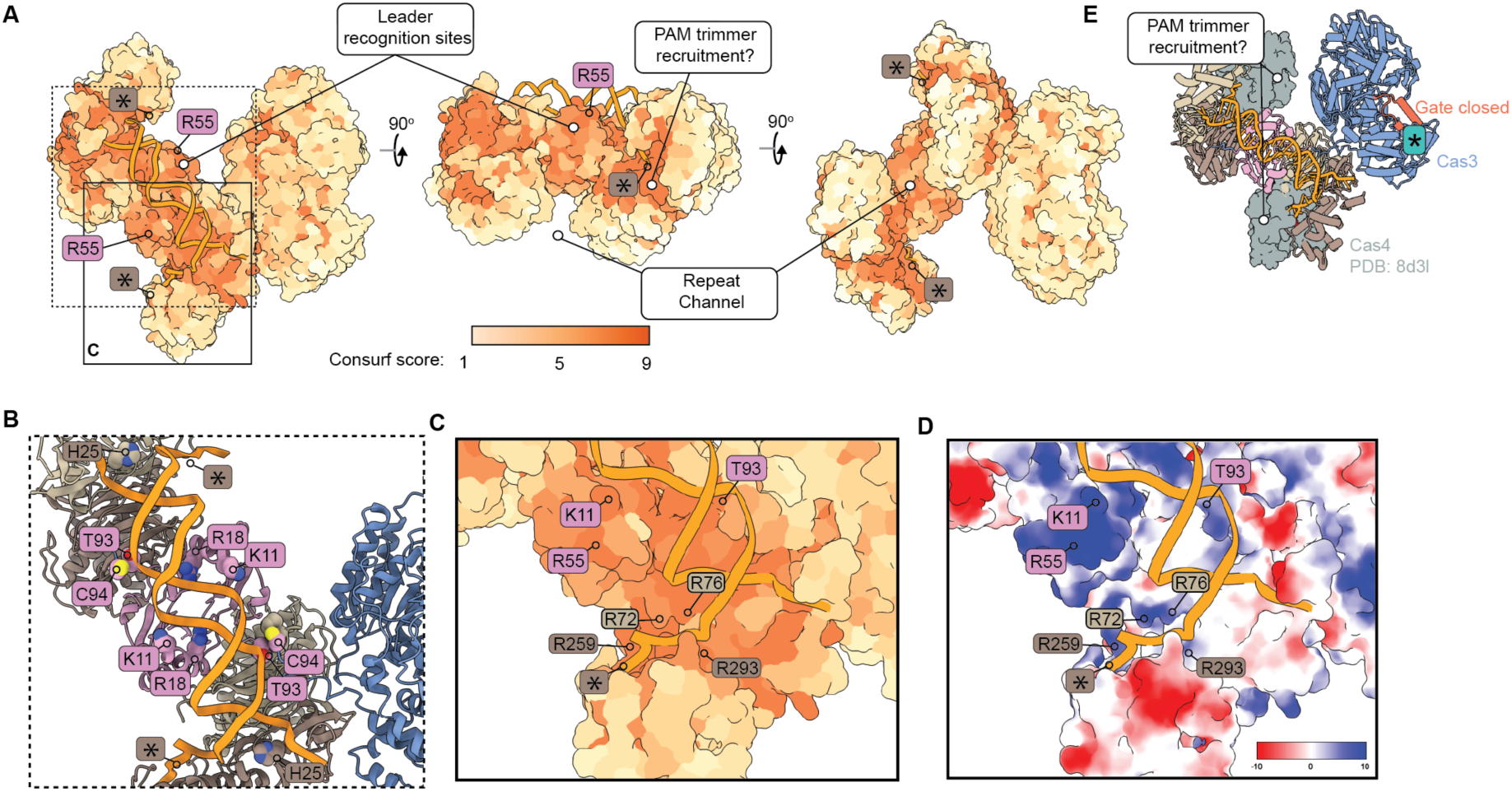
3’ ssDNA channel and additional DNA binding sites are conserved. Related to Figure 3. (A) Multiple sequence alignments of Cas1 (n=679) and Cas2/3 (n=745) from type I-F systems were used to calculate Consurf conservation scores, that are mapped back to the atomic model of the Cas1-2/3 complex bound to foreign DNA (1 – least conserved is light yellow, while 9 – most conserved is dark orange). Conserved residues around the Cas1 active site are positioned to facilitate recruitment of a PAM-trimming nuclease. Asterisk (*) indicates the Cas1 transesterification sites. (B) Cas2 residues hydrogen bond with the DNA backbone to position the DNA fragment symmetrically in the foreign DNA binding channel. Asterisk (*) indicates the Cas1 transesterification sites. (C-D) The electrostatic funnel that positions the 3’ end of the foreign DNA in the Cas1 active site is conserved. Atomic model in D colored by electrostatic potential (kcal/mol\**e* at 298 K) in ChimeraX. (E) DNA-binding triggered conformational changes position the HD domain away from the PAM but expose faces that recruit Cas4 in other CRISPR systems. The RecA1 gate of the HD domain (colored orange) remains in the closed position after foreign DNA binding. A previously determined structure Cas1-Cas2-Cas4 (PDB: 8d3l) was docked into the foreign DNA-bound structure using the mmaker command in ChimeraX. The Cas1 and Cas2 from 8d3l were hidden and Cas4 is displayed in grey with surface representation. Asterisk (*) indicates the HD domain active site of Cas3.

**Figure S10.**
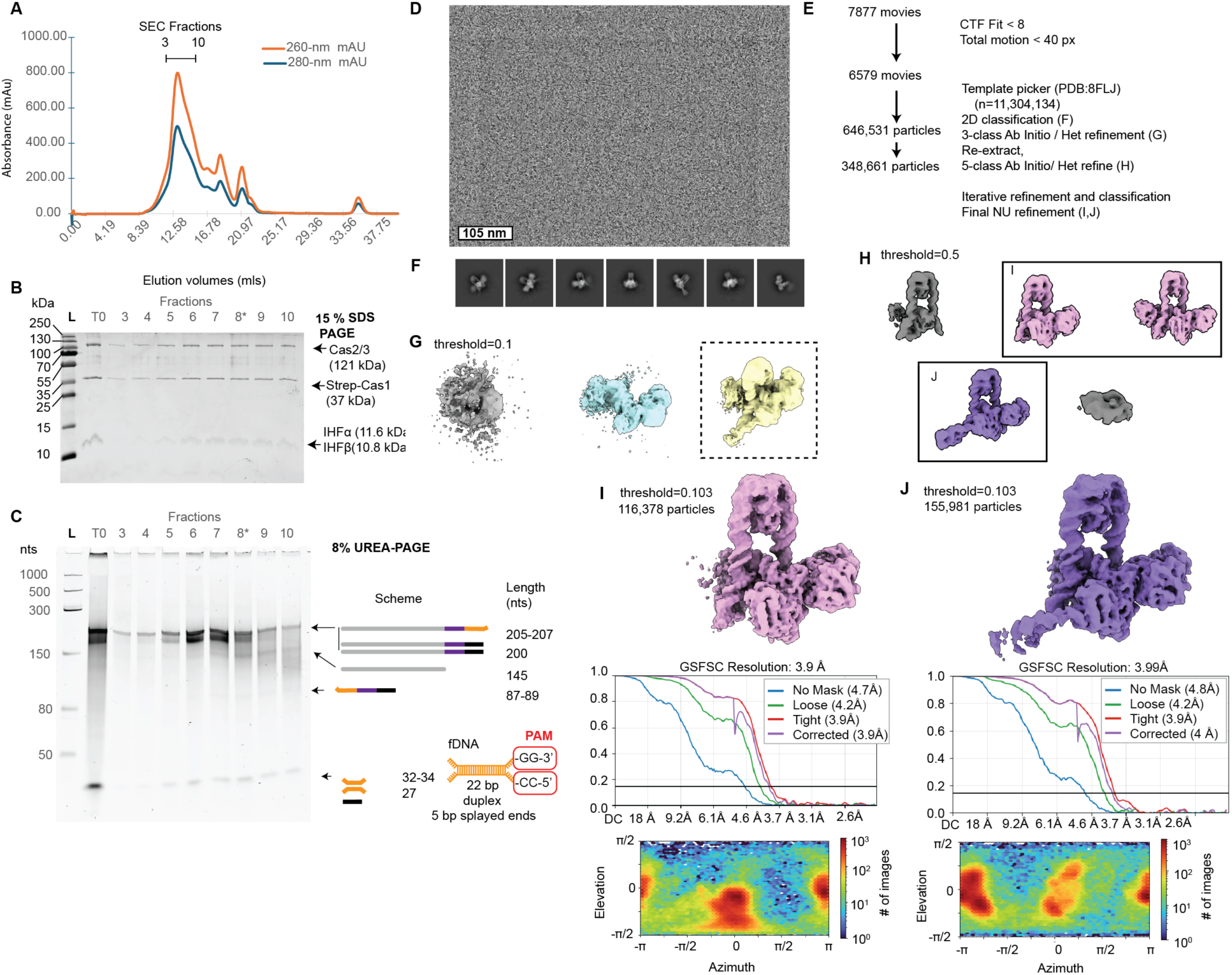
Purification and cryo-EM data analysis of PAM-containing integration complex. Related to Figure 4. (A) Size exclusion chromatogram of the integration reaction after 10 minutes at 25 °C. (B) 15 % SDS PAGE gel of the main peak fractions reveals all of the protein components of the integration complex are present. Fraction 8 (green box) was frozen for cryo-EM. (C) 8 % Urea-PAGE gel of main peak fractions reveals that all of the DNA components of a partial integration reaction are present in fraction 8. (D) Sample micrograph from data collection. (E) Data analysis workflow. (F) Representative 2D class averages selected for further processing. (G) 3-class *ab initio* and Heterogeneous refinement identifies a single integration complex class (black box). (H) Five-class ab-initio reconstruction and heterogeneous refinement of integration complex particles. Black boxes indicate particle stacks used in subsequent analyses (I) Non-uniform refinement of 116,378 particles containing the U-bend but not the loop indicative of the first transesterification, FSC curve and azimuth plot displayed below (threshold 0.103). (J) Non-uniform refinement of 155,981 particles containing the U-bend and loop indicative of the first transesterification FSC curve and azimuth plot displayed below (threshold 0.103).

**Figure S11.**
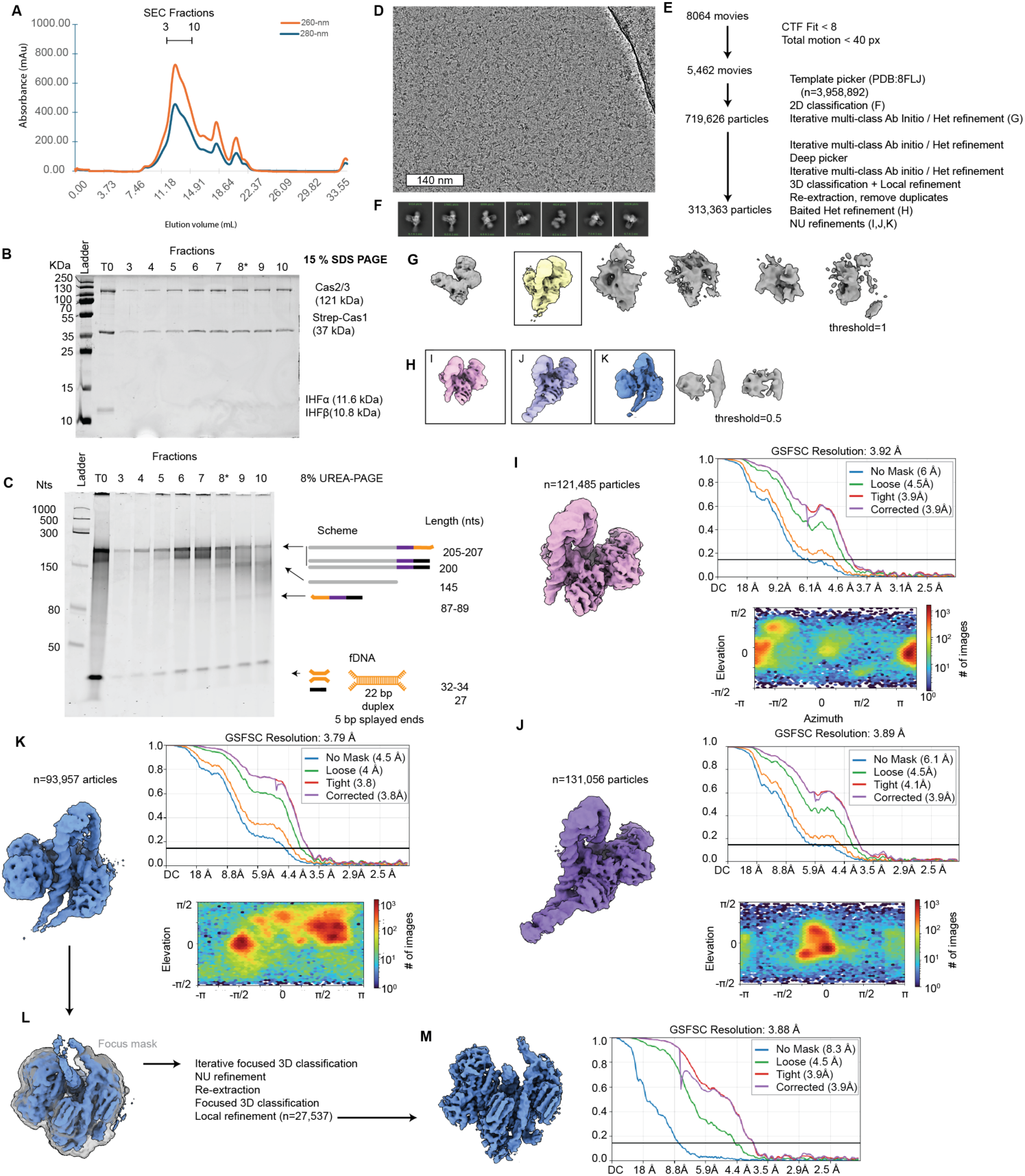
Purification and cryo-EM data analysis of PAM-containing integration complex. Related to Figure 4 and Figure 5. (A) Size exclusion chromatogram of the integration reaction after 10 minutes at 25 °C. (B) 15 % SDS PAGE gel of the main peak fractions reveals all of the protein components of the integration complex are present. Fraction 8, marked with an asterick (*) was frozen for cryo-EM. (C) 8 % UREA-PAGE gel of main peak fractions reveals that all of the DNA components of a partial integration reaction are present in fraction 8. (D) Sample micrograph from data collection. (E) Data analysis workflow. (F) Representative 2D class averages selected for further processing. (G) 6-class *ab initio* and heterogeneous refinement identifies a single integration complex class (black box). (H) 6-class *ab initio* and heterogeneous refinement of the integration complex class from panel G identifies multiple distinct complexes (black outlines). (I) Non-uniform refinement, FSC curve and azimuth plot for the integration complex prior to the first integration reaction. (J) Non-uniform refinement, FSC curve and azimuth plot for the integration complex after the first integration reaction, which contains the DNA loop that is directed by IHF into the first transesterification site. (K) Non-uniform refinement, FSC curve and azimuth plot for the integration complex after the second integration reaction, when both strands of the repeat have been covalently linked to the foreign DNA.

**Figure S12.**
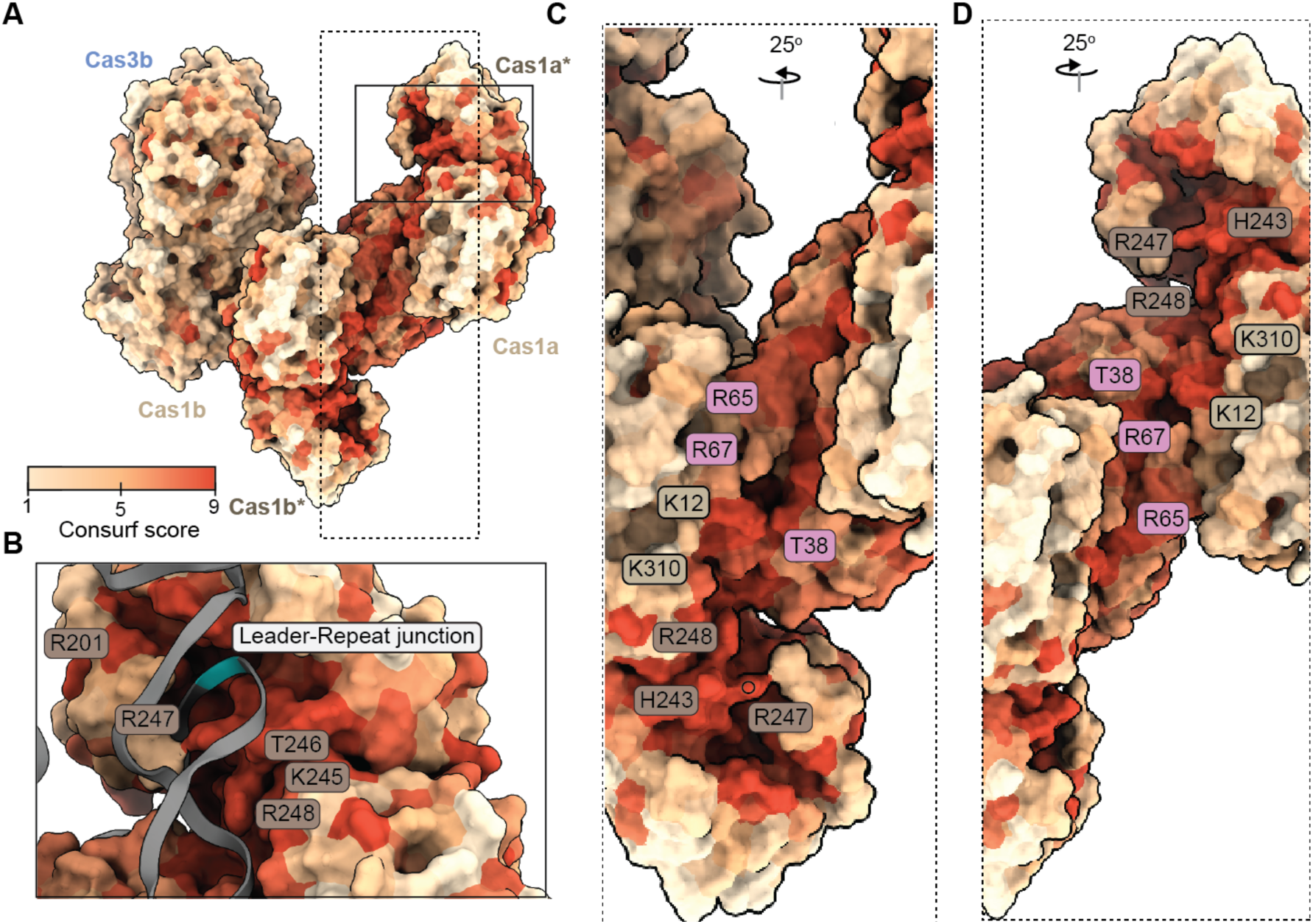
Positively charged residues in the repeat channel are conserved in type I-F CRISPR systems. Related to Figure 4 and Figure 5. (A) Consurf analysis of the integration complex with DNA removed to show the conserved residues of the repeat channel. (B) K245, T246, R247, and R248 are conserved and interact with the B-form portion of the CRISPR repeat in the channel just after the leader-repeat junction. (C, D) Conserved basic and polar residues line the region of the repeat channel that distorts the repeat DNA.

**Figure S13.**
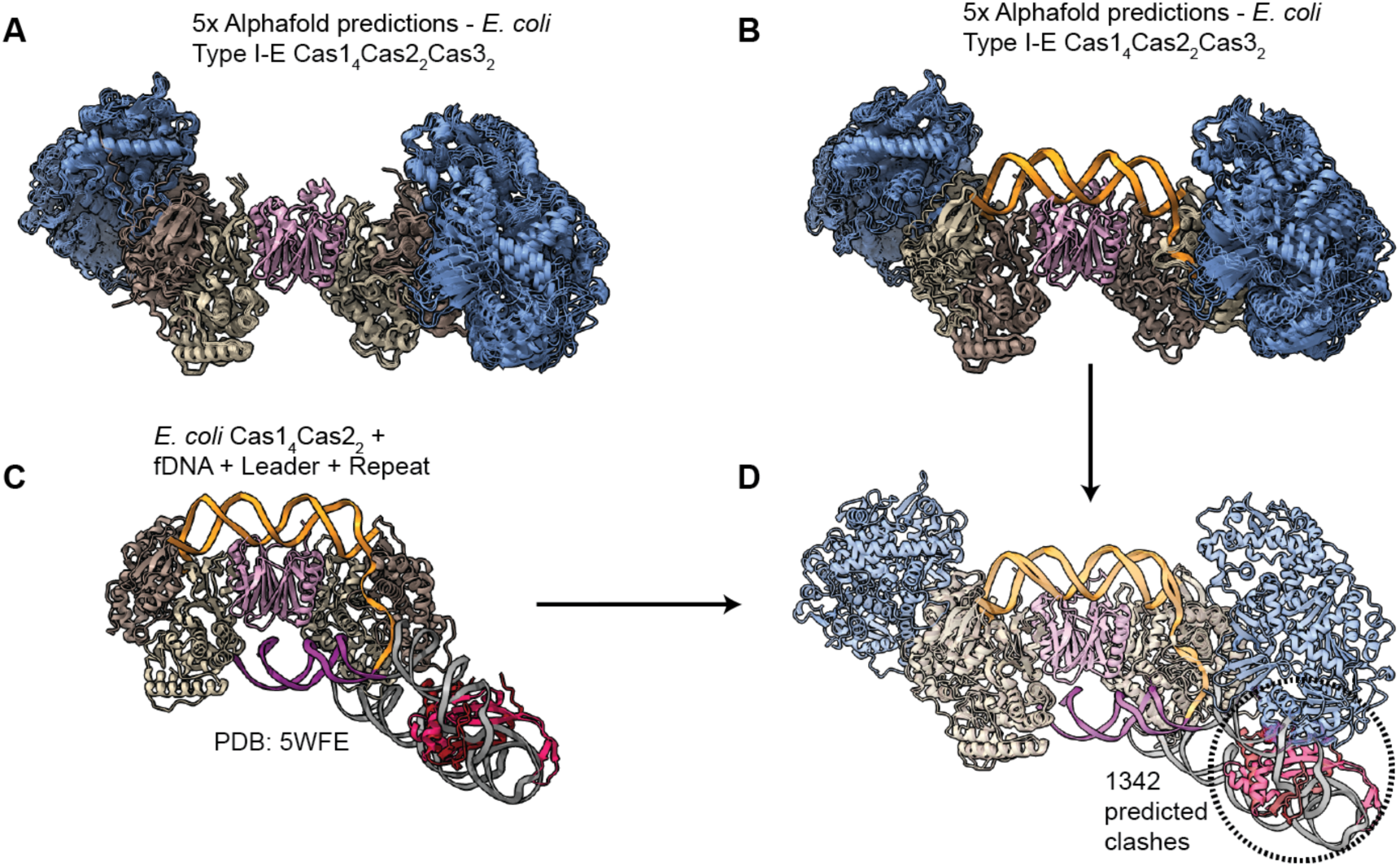
Cas3 in a predicted Cas1-2-3 complex from *E. coli* blocks interactions between the folded genome and Cas1. Related to Figure 1, Figure 3, Figure 4. (A) All five predicted complexes from an AlphaFold3 prediction of a Cas1_4_Cas2_2_Cas3_2_ hetero-octamer predict Cas3 in the same conformation on the Cas1 dimer (mean iPTM across five models= 0.48). (B) Addition of a short dsDNA fragment to the prediction does not change the location of the predicted Cas1-Cas3 association (mean iPTM across five models = 0.66). (C) The Cas1-2 integration complex from *E. coli* docked onto an AlphaFold3 predicted structure of the DNA-bound Cas1_4_Cas2_2_Cas3_2_ hetero-octamer reveals 1342 atomic clashes between the folded leader, which interacts with Cas1, and Cas3, which is predicted to interact with Cas1 at the same site.

## SUPPLEMENTAL TABLES

**Table S1. Refinement statistics for all models and maps**

**Table S2. Table of Cas2 RMSDs and buried surface area at homodimer interface**

